# The maternal microbiome modifies adverse effects of protein undernutrition on offspring neurobehavioral impairment in mice

**DOI:** 10.1101/2024.02.22.581439

**Authors:** Elena J. Coley-O’Rourke, Gregory R. Lum, Geoffrey N. Pronovost, Ezgi Özcan, Kristie B. Yu, Janet McDermott, Anna Chakhoyan, Eliza Goldman, Helen E. Vuong, Jorge Paramo, Alison Chu, Kara L. Calkins, Elaine Y. Hsiao

## Abstract

Protein undernutrition is a global risk factor for impaired growth and neurobehavioral development in children. However, the critical periods, environmental interactions, and maternal versus neonatal influences on programming lasting behavioral abnormalities are poorly understood. In a mouse model of fetal growth restriction, limiting maternal protein intake particularly during pregnancy leads to cognitive and anxiety-like behavioral abnormalities in adult offspring, indicating a critical role for the gestational period. By cross-fostering newborn mice to dams previously exposed to either low protein or standard diet, we find that the adult behavioral impairments require diet-induced conditioning of both fetal development and maternal peripartum physiology, rather than either alone. This suggests that protein undernutrition during pregnancy directly disrupts fetal neurodevelopment and indirectly alters maternal state in ways that interact postnatally to precipitate behavioral deficits. Consistent with this, maternal protein restriction during pregnancy reduces the diversity of the maternal gut microbiome, modulates maternal serum metabolomic profiles, and yields widespread alterations in fetal brain transcriptomic and metabolomic profiles, including subsets of microbiome-dependent metabolites. Depletion of the maternal microbiome in protein-restricted dams further alters fetal brain gene expression and exacerbates neurocognitive behavior in adult offspring, suggesting that the maternal microbiome modifies the impact of gestational protein undernutrition on risk for neurobehavioral impairment in the offspring. To explore the potential for microbiome-targeted interventions, we find that maternal treatment with short chain fatty acids or a cocktail of 10 diet- and microbiome-dependent metabolites each yield differential effects on fetal development and/or postnatal behavior. Results from this study highlight impactful prenatal influences of maternal protein undernutrition on fetal neurodevelopment and adverse neurobehavioral trajectories in offspring, which are mitigated by microbiome-targeted interventions during pregnancy.

## Main

Protein undernutrition is a global risk factor for childhood stunting^1^, which is co-morbid with lasting neurological disabilities, including cognitive impairment and anxiety^2–4^. In humans and animal models, abnormalities in the maturation and function of the gut microbiome contribute to malnutrition-induced stunting^5^, but standard therapeutic foods have limited effectiveness in supporting persistent microbial rehabilitation^5–7^. There is increasing evidence that bacterial treatments and custom microbiota-directed diets ameliorate growth restriction in animal models of malnutrition and in malnourished children^8–11^. This raises the prospect of using microbiome-based treatments to combat malnutrition-induced growth defects. However, current medical and nutritional interventions that treat childhood stunting are often inadequate to ameliorate co-morbid neurobehavioral impairments^12^. Individuals who experienced protein-energy malnutrition during the first year of life displayed cognitive impairment and depressive symptoms during adolescence and adulthood, despite adequate nutritional rehabilitation and growth recovery during childhood^13–16^. Similarly, supplementing stunted infants with a milk-based formula failed to ameliorate heightened anxiety and cognitive impairments in adolescence^3,4^. Whether alterations in the microbiome contribute to the neurological comorbidities caused by protein undernutrition, and whether microbiome-based interventions can be used to ameliorate them, is poorly understood.

Brain abnormalities, such as cerebral atrophy, ventricular dilation, and myelin-related deficits, are seen in protein malnourished infants as young as 3 months of age and through 36 months of age^17,18^, highlighting an early critical period during which protein undernutrition impairs neurodevelopment. In animal models, restricting protein intake particularly during pregnancy yields persistent neurological and neurobehavioral impairments in the offspring, including abnormalities in neuronal proliferation and apoptosis^19^, neocortical activity^20^, hippocampal morphology^21^, learning and memory, and anxiety-like behaviors^19–22^. These results indicate that adverse neurological consequences of protein undernutrition can originate from gestational influences^23^. In considering potential contributions of the microbiome, evidence from animal models indicate that alterations in the maternal microbiome contribute to adverse effects of immune activation, stress, and high-fat diet on neurological and behavioral deficits in the offspring^24–26^, either by directing fetal neurodevelopment during pregnancy^27,28^ or by shaping early postnatal neural development via vertical transmission at birth and postpartum^28,29^. These findings align with human studies reporting that alterations in the maternal microbiome during pregnancy are associated with abnormalities in offspring behavior^30,31^ and that postpartum nursing supports cognitive, language, and microbiome development in the first years of life^32^. However, mechanisms by which maternal protein undernutrition leads to lasting neurobehavioral deficits in the offspring, and how these processes may be modified by the microbiome, are unknown.

Herein, we examine effects of maternal protein undernutrition during pregnancy on maternal-fetal health and offspring behavior in mice. We use a cross-fostering paradigm to evaluate the ability of maternal protein restriction to directly alter fetal neurodevelopment and indirectly condition maternal physiology to engender lasting cognitive and anxiety-related behaviors in adult offspring. We profile effects of maternal protein restriction on the maternal microbiome, and further assess the impact of maternal microbiome depletion and microbial metabolite supplementation on maternal-fetal and offspring behavioral responses to maternal protein restriction. Results from this study highlight the importance of the gestational period in protein undernutrition altering both maternal health and fetal developmental trajectories. Furthermore, results reveal a role for the maternal microbiome in modulating the severity of fetal developmental and adult neurobehavioral impairments induced by maternal protein undernutrition. These advances in illuminating molecular underpinnings of adverse behavioral outcomes of protein undernutrition could potentially lead to new approaches to ameliorate neurological disorders that co-occur with impaired growth in malnourished children.

## Results

### Maternal protein restriction yields behavioral abnormalities in adult offspring by altering both fetal development and maternal postpartum physiology

Protein undernutrition is associated with both childhood stunting and neurological dysfunction, but the etiopathogenesis of lasting neurobehavioral deficits remains poorly understood. To explore this relationship, we first evaluated effects of maternal protein restriction particularly during pregnancy on behavior in adult offspring. To model maternal protein undernutrition, male and female C57Bl/6J mice were fed a 6% protein diet (protein restriction, PR) or 20% protein control diet (CD) for 2 weeks prior to timed-mating and throughout gestation (**Fig. 1a**). PR and CD formulations were isocaloric, where the 14% protein content lacking in PR was replaced by carbohydrates, mainly sucrose and cellulose (**Extended Data Fig. 1a, Supplementary Table 1**). Consistent with existing literature using this paradigm as a model of fetal growth restriction (FGR)^33,34^, maternal consumption of a PR diet prior to and throughout pregnancy resulted in reduced fetal weight at late gestation (embryonic day (E) 18.5) and maternal net weight loss with no difference in diet consumption, in addition to elevated maternal serum corticosterone and reduced litter size at birth (**Extended Data Fig. 1b-f**). These findings suggest a state of maternal physiological stress.

**Figure 1:**
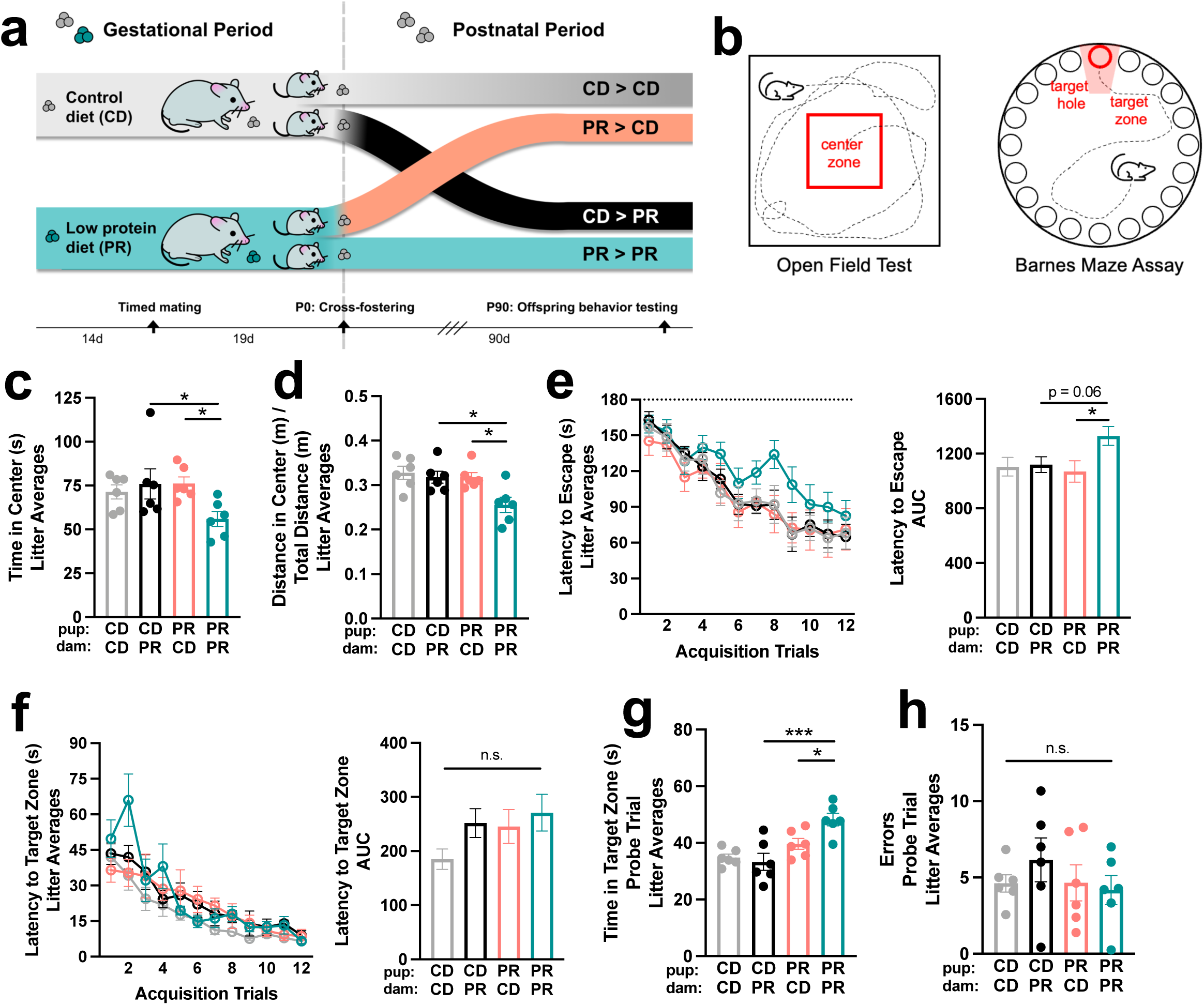
The prenatal period of maternal protein restriction is critical for programming cognitive and anxiety-like behavioral deficits in adult offspring. a, Graphic of cross-fostering paradigm. b, Graphic of behavioral tests, open field test (left), Barnes maze (right). c, Time in center in open field test, all offspring averaged within each litter (two-way ANOVA with Sidak, n = 6 litters per group). Top row refers to pup condition, bottom row refers to dam condition. d, Distance in center in open field test, controlled by total distance traveled, all offspring averaged within each litter (two-way ANOVA with Sidak, n = 6 litters per group). Top row refers to pup condition, bottom row refers to dam condition. e, Left: Latency to escape in Barnes maze acquisition phase, all offspring averaged within each litter (n = 6 litters per group). Right: AUC of latency to escape (two-way ANOVA with Sidak). Top row refers to pup condition, bottom row refers to dam condition. f, Left: Latency to target zone in Barnes maze acquisition phase, all offspring averaged within each litter (n = 6 litters per group). Right: AUC of latency to target zone (two-way ANOVA with Sidak). Top row refers to pup condition, bottom row refers to dam condition. g, Time in target zone in Barnes maze probe trial, all offspring averaged within each litter (two-way ANOVA with Sidak, n = 6 litters per group). h, Errors made in Barnes maze probe trial, all offspring averaged within each litter (two-way ANOVA with Sidak, n = 6 litters per group). Mean +/- SEM, *p < 0.05, **p < 0.01, ***p < 0.001, ****p < 0.0001, n.s. not significant.

To decouple lasting effects of maternal protein restriction during pregnancy on maternal postpartum physiology versus fetal neurodevelopment, pups were cross fostered at birth to dams gestationally exposed to PR or CD to form the following groups: CD pups fostered to CD dams (CD è CD), PR pups fostered to CD dams (PR è CD), CD pups fostered to PR dams (CD è PR), PR pups fostered to PR dams (PR è PR) (**Fig. 1a**). Moreover, to examine effects of maternal protein undernutrition, specifically during pregnancy, on postnatal health of the offspring, PR-fed dams were switched to CD at parturition, and all pups were reared on CD from birth in order to isolate PR to the gestational period. Of all the experimental groups, PR pups fostered to PR dams exhibited the smallest litter sizes by weaning age (**Extended Data Fig. 1g**) and the lowest total litter survival at 26.5% (**Extended Data Fig. 1h**). CD pups fostered to PR dams exhibited intermediate reductions in litter survival to 46.7%, whereas PR pups fostered to CD dams exhibited 100% survival, as did CD pups fostered to CD dams. These results indicate that offspring survival is determined by an interaction between direct effects of gestational PR on fetal health and indirect effects of gestational PR on maternal health persisting into the postnatal period. It further indicates that gestational PR-induced alterations in maternal, but not fetal health, are sufficient to reduce offspring survival. There were no significant differences in pup weights in the first two weeks of life (**Extended Data Fig. 1i**), suggesting that fetuses from PR dams exhibit weight recovery with postnatal rearing on CD.

To assess lasting effects of maternal protein restriction during pregnancy on adverse neurobehavioral outcomes in the offspring, fostered offspring were weaned, reared to adulthood, and tested in behavioral assays related to anxiety and cognition (**Fig. 1b**). The open field assay is a benchmark test for stress-induced thigmotaxis as an endophenotype of anxiety-like behavior^35,36^. Adult PR offspring previously fostered to PR dams and reared since birth on CD (PRèPR) displayed significantly reduced time and distance in the center of open field (**Fig. 1c-d**), as compared to other experimental groups, with no difference in average speed or total distance traveled (**Extended Data Fig. 2a**). This anxiety-like phenotype was more striking in females than males, with postnatal influence of CD rearing appearing to be sex-discriminatory (**Extended Data Fig. 2b**). This female-bias in anxiety-like response was similarly reported in a multi-hit pre- and postnatal adversity model^37^. This finding suggests that adverse postnatal interactions between maternal and fetal responses to gestational PR lead to anxiety-like behavior. The Barnes maze test is a benchmark assay of spatial learning and memory^38^, wherein mice are trained over repeated trials to identify which out of 20 holes contains an escape box (acquisition phase) and their ability to recall the spatial location of the escape is tested in a final probe trial 24 hours post-training. During the acquisition phase, adult PRèPR offspring exhibited increased latency to escape (**Fig. 1e**), but no significant difference in primary latency to target zone (**Fig. 1f**), suggesting deficient learning. During the probe phase, these PRèPR offspring displayed increased time in the target zone (**Fig. 1g**), but no significant difference in errors made (**Fig. 1h**), suggesting adequate 24-hour recall, but increased perseveration. There were no overt sex differences in cognitive deficits seen in offspring of PR-fed dams (**Extended Data Fig. 2c-f**). These findings suggest that maternal PR during pregnancy alters both maternal physiology and fetal development in ways that interact postnatally to engender lasting behavioral impairments in offspring with adequate postnatal nutrition.

### Maternal protein restriction alters functional signatures in the fetal brain

Maternal malnutrition can alter the trajectory of fetal neurodevelopment to result in long-term changes in brain function and behavior^19–22^. To uncover mechanisms by which maternal protein undernutrition programs adverse behavioral outcomes in the offspring, we first examined the effects of maternal PR on E18.5 fetal brains by transcriptomic profiling. We identified 505 significantly up-regulated and 423 significantly down-regulated genes in response to maternal PR (**Fig. 2a, Supplementary Table 2**). By gene ontology analysis, the upregulated genes related to biological processes such as neuronal apoptosis (reported to be increased in fetal brain following protein restriction^19^), ion and amino acid transport, and response to growth factor (**Fig. 2b**), suggesting compensatory mechanisms related to brain sparing^33,39^. Downregulated genes related to biological processes such as T-cell differentiation and nervous system development potentially highlight immune or neuroimmune disruptions, as previously implicated in malnourished children^40^. Particular neurobehaviorally-relevant genes were significantly altered in fetal brains from PR dams, including downregulated serotonin transporter (*Slc64a)* and serotonin receptor (*Htr3a)*, insulin-like growth factor 1 (*Igf1*), which modulates anxiety and stress response^41^, and Ataxin 1 (*Atxn1*), related to learning and hippocampal deficits^42^. Conversely, activity-dependent neuroprotective protein (*Adnp*), implicated in cognitive and social impairments and stress response^43^, and dopamine beta hydroxylase (*Dbh*), relevant to learning and memory^44^ were upregulated in fetal brains from PR dams (**Supplementary Table 2).**

**Figure 2:**
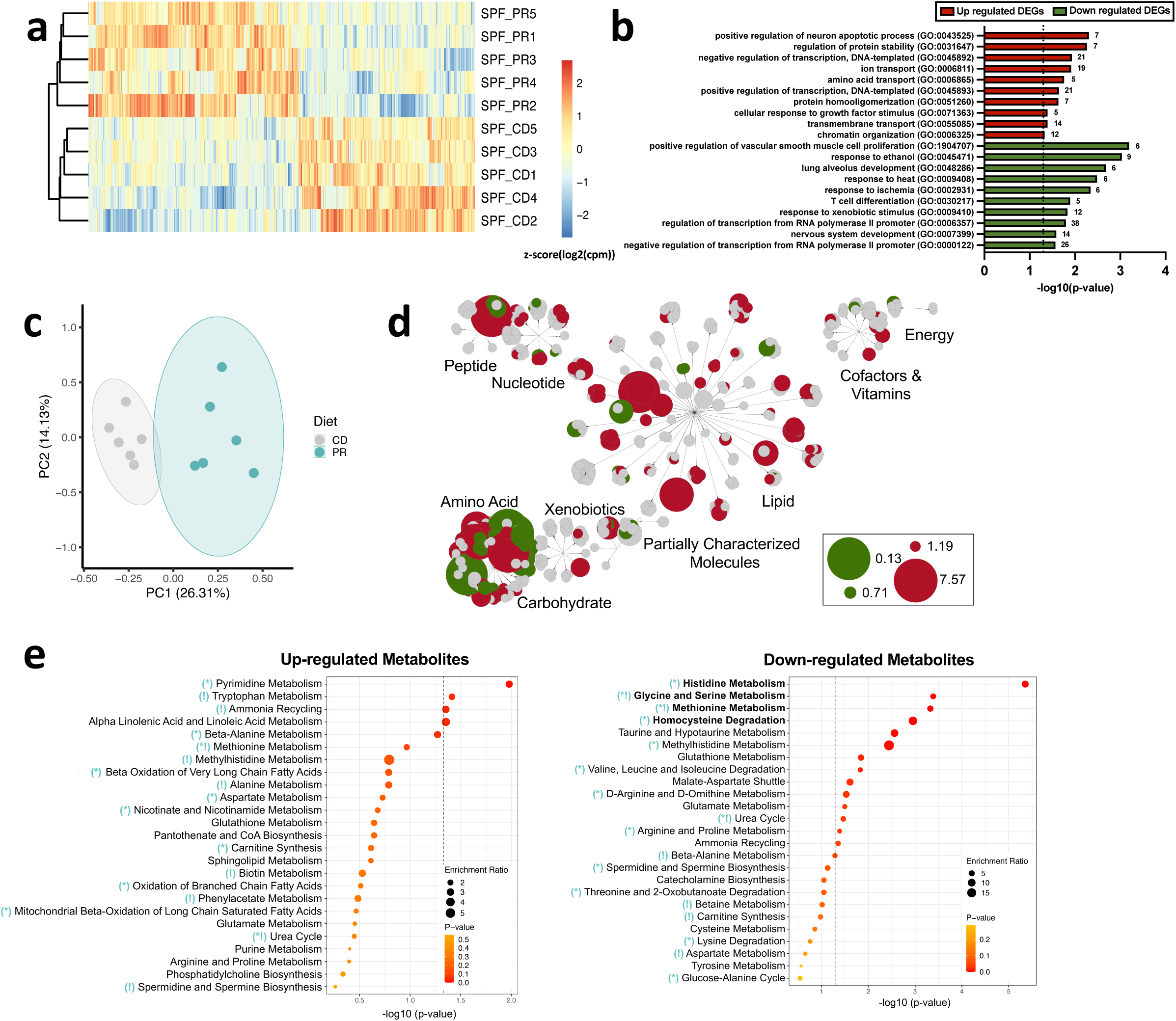
Maternal protein restriction alters transcriptional and metabolomic profiles in the fetal brain. **a,** Heatmap of differentially expressed genes (DEGs) (505 up-regulated, 423 down-regulated; p < 0.05) of E18.5 fetal brains from SPF CD or SPF PR dams with Euclidean clustering on rows (n = 5 litters per group, 2 brains pooled per litter). **b,** Biological Process gene ontology (GO) of upregulated DEGs (p < 0.05, red) and downregulated DEGs (p < 0.05, green) of transcripts from E18.5 fetal brains from SPF CD or SPF PR dams (n = 5 litters per group, 2 brains pooled per litter). **c,** PCA of untargeted metabolomics from E18.5 fetal brains from SPF CD and SPF PR dams (n= 6 litters per group, 1.5-2 brains pooled per litter). **d,** Metabolon network map showing positive and negative fold change. **e,** Top enriched Small Molecule Pathway Database pathways for significantly upregulated metabolites (p < 0.05, left) and downregulated metabolites (p < 0.05, right) for E18.5 fetal brains from SPF PR dams compared to SPF CD dams (bold pathways have q < 0.05; teal symbols relate to analogous enriched pathways in SPF PR vs SPF CD maternal serum [see Extended Data Fig. 3c]; * = enriched pathway in same direction; ! = enriched pathway in opposite direction).

These PR-induced alterations in fetal gene expression corresponded with widespread changes in levels of fetal brain metabolites. Liquid chromatography tandem mass spectrometry-based metabolomic profiling of E18.5 brains yielded detectable levels of 681 identified compounds, spanning amino acid, peptide, carbohydrate, energy, lipid, nucleotide, cofactor and vitamin, and xenobiotic super pathways. By principal component analysis, metabolomic profiles of fetal brains from PR dams were clearly discriminated from CD controls along PC1 (**Fig. 2c**), with statistically significant alterations in 220 metabolites (**Supplementary Table 3**). The most highly affected metabolite classes were amino acids (98) and lipids (84) (**Fig. 2d**). Of the amino acid-related metabolites that were significantly altered in fetal brains from PR dams relative to CD controls, 50% were increased (and 50% were decreased), suggesting complex regulation of amino acids. By comparison, we further profiled metabolomics in maternal serum and found 332 serum metabolites that were statistically significantly altered by PR consumption (**Extended Data Fig. 3a**, **Supplementary Table 4**). As in fetal brains, the majority were amino acids (107) and lipids (144) (**Extended Data Fig. 3b**). However, in contrast to fetal brains, the alterations in amino acid-related metabolites seen in maternal serum were mostly reductions (81.3%, as compared to 50% in fetal brain) (**Supplementary Table 3, 4**), which is consistent with low protein intake and evidence of fetal brain sparing^39^. In line with this, all essential amino acids and glucose showed significant reductions in PR maternal serum compared to CD (**Extended Data Fig. 3d-m**). However, glucose, lysine, and phenylalanine showed no significant difference between PR and CD fetal brain, and tryptophan showed a significant increase in PR fetal brain compared to CD, further supporting a metabolite-specific brain-sparing phenotype. (**Extended Data Fig. 3d**, h, j, l).

By metabolite set enrichment analysis, fetal brain metabolites that were reduced by maternal PR mapped to 14 metabolic pathways, with histidine metabolism, glycine and serine metabolism, methionine metabolism, and homocysteine degradation persisting after statistical correction (**Fig. 2e**). The fetal brain metabolites that were increased by maternal PR mapped to 4 pathways, including pyrimidine metabolism and tryptophan metabolism, although none survived statistical correction (**Fig. 2e**). Similar to fetal brain metabolites, maternal serum metabolites related to pyrimidine metabolism were upregulated by PR, whereas pathways related to tryptophan metabolism and valine, leucine, and isoleucine degradation were downregulated (**Extended Data Fig. 3c**). The most severely depleted individual metabolites in PR fetal brains included urea (10.1% the levels of CD fetal brains), a product of protein catabolism, and hypotaurine (24.0% the levels of CD fetal brains), a precursor of taurine, which directly supports synaptic formation and transmission in the developing brain^45^. Conversely, PR fetal brains had a 276.9% increase in corticosterone, which is consistent with increased corticosterone in maternal serum by ELISA (**Extended Data Fig. 1e**). Synthetic corticosteroids which mimic glucocorticoid activity during gestation have been previously reported to impair hippocampal synaptic development in primates^46^, and myelination and brain growth in sheep^47,48^. Furthermore, maternal cortisol has been correlated with sex-specific amygdala connectivity and internalizing symptoms in humans^49^. Taken together, these results indicate that maternal protein undernutrition during pregnancy modifies transcriptional and metabolomic profiles in the late gestational fetal brain, which can contribute to fetal neurodevelopmental programming in ways that interact postnatally with maternal factors to yield behavioral abnormalities in adult offspring.

### Murine maternal protein restriction and human fetal growth restriction are associated with reduced diversity of the maternal and infant gut microbiome

Maternal behavioral and physiological care during early life has profound and persistent effects on offspring neurobehavioral development^50,51^. Results from our cross-fostering experiments indicate that PR-induced alterations in offspring behavior require antenatal exposure of both fetuses and mothers to PR during gestation (**Fig. 1**). To gain insight into how protein undernutrition during pregnancy impacts maternal health to disrupt offspring behavioral development, we first tested dams for postpartum behavior related to maternal care. The pup retrieval test is a benchmark behavioral assay that assesses the mother’s response to retrieve pups upon their removal from the nest^52,53^. There were no statistically significant differences between PR and CD dams in their latency to full retrieval of their fostered pups, either PR or CD (**Extended Data Fig. 1j**). Consistent with this, there were no differences in pup righting reflex or ultrasonic vocalizations upon maternal separation (**Extended Data Fig. 1k-l**). These results suggest no overt difference in pup-directed maternal care across dams previously fed PR or CD during pregnancy.

In addition to maternal care, the maternal gut microbiome is increasingly appreciated for its roles in promoting healthy neonatal development by supporting nutrition and immune development^23,54^. Alterations in the maternal microbiome, observed in various models of malnutrition^55,56^, can disrupt early neurodevelopment and elicit lasting changes in offspring behavior^24,25,27,28,30,31,57,58^. To determine effects of protein restriction on the composition of the maternal gut microbiota, we performed 16S rRNA gene sequencing of fecal samples collected over the course of gestation from dams fed PR or CD (**Supplementary Table 5**). Consistent with previous reports^59^ bacterial alpha-diversity increased with pregnancy in CD-fed dams (**Fig. 3a**). However, this was prevented by PR, as fecal microbiota from PR dams displayed significantly decreased Shannon diversity [a measure of both richness (the number of different species) and evenness (the amount within each species)] and Pielou’s evenness as compared to CD controls (**Fig. 3a**). Principal coordinates analysis of weighed Unifrac distances discriminates microbiota from PR dams away from CD controls by E10.5 (**Fig. 3b**). Differences in beta diversity were driven by statistically significant alterations in the relative abundances of 31 bacterial taxa (**Fig. 3c**), which were dominated by *Clostridia* species. Reductions align with the finding that *Clostridia* are the main metabolizers of dietary protein and amino acids^60^. Altogether, these results indicate that maternal protein undernutrition during pregnancy reduces diversity of the maternal microbiota, which corresponds with alterations in functional profiles in the fetal brain and neurobehavioral deficits in adult offspring despite being reared on a standard protein diet since birth.

**Figure 3:**
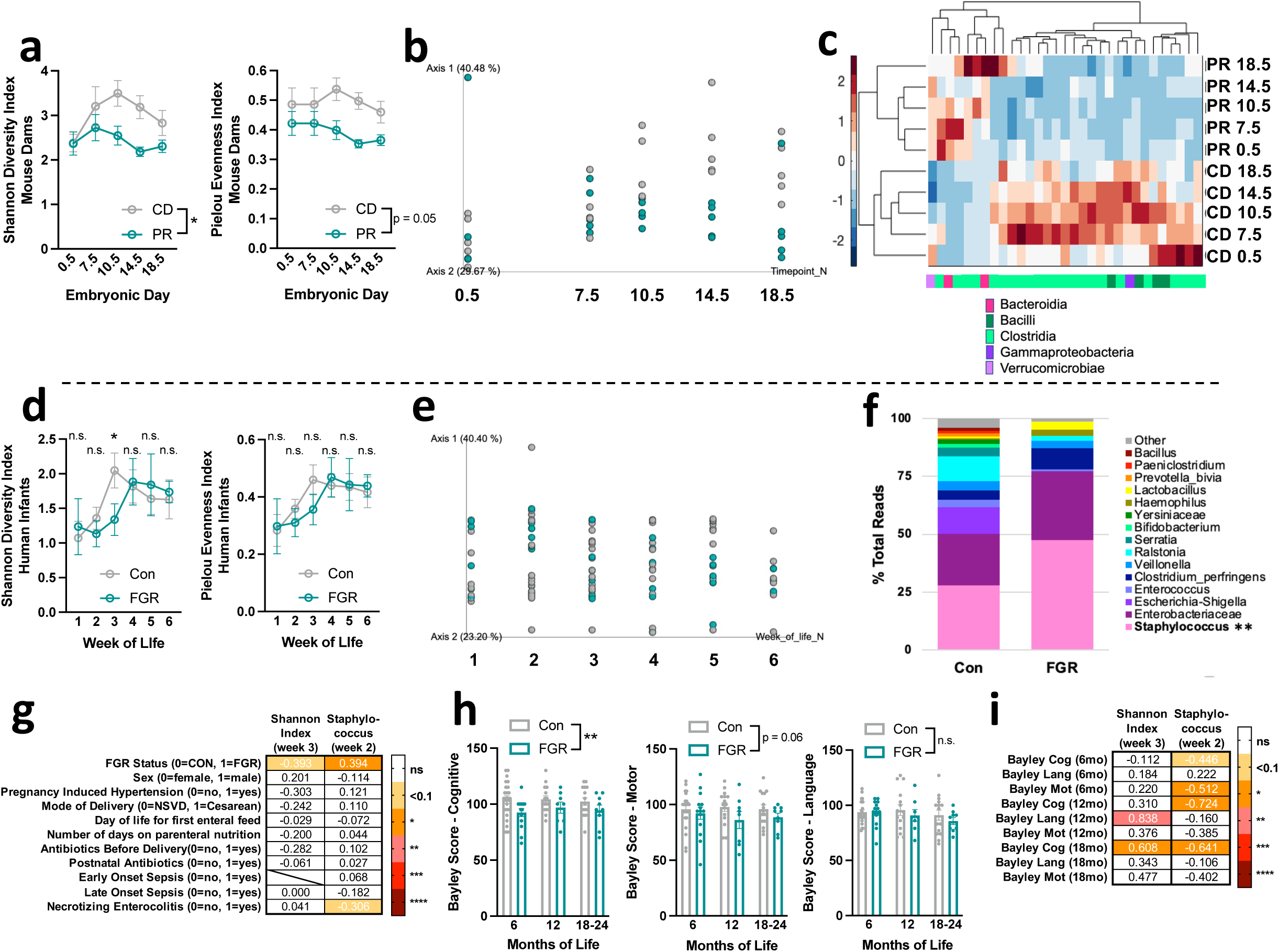
Murine maternal protein restriction or human fetal growth restriction reduces diversity of the gut microbiome. **a,** Left: Shannon diversity index across gestation in SPF CD and SPF PR dams (two-way repeated measures ANOVA with Sidak, n = 5 dams per group). Right: Pielou evenness index across gestation in SPF CD and SPF PR dams (two-way repeated measures ANOVA with Sidak, n = 5 dams per group). **b,** PCA of weighted unifrac across gestation in SPF CD and SPF PR dams (n = 5 dams per group). Axis 3 = time pseudo-axis. **c,** Heatmap of significantly different taxa across gestation in SPF CD and SPF PR dams (Qiime2 Kruskal Wallis test, q < 0.05, n = 5 dams per group). **d,** Left: Shannon diversity index in the first 6 weeks of life in Con and FGR infants (unpaired Welch’s t-test per timepoint, n = 37 (with 1-4 timepoints each), 16 (with 1-5 timepoints each) infants from top to bottom). Right: Pielou evenness index in the first 6 weeks of life in Con and FGR infants (unpaired Welch’s t-test per timepoint, n = 37 (with 1-4 timepoints each), 16 (with 1-5 timepoints each) infants from top to bottom). **e,** PCA of weighted unifrac of Con and FGR infants across the first six weeks of life (n = 34 Con with 1-4 timepoints each, 16 IUGR with 1-5 timepoints each). **f,** Taxa bar plots of taxa >1% total reads in Con or FGR infants in week 2 of life (n = 26, 10, infants from left to right, ** survived correction by Kruskal Wallis test in Qiime2). **g,** Spearman correlation matrix of Shannon diversity index at week 3 and Staphylococcus genus % reads at week 2, against clinical metadata (Con n = 16, 22; FGR n = 9, 10, from left to right). Displayed values are R^2^. **h,** Left: Bayley Cognitive assessment scores across the first two years of life in Con and FGR infants (2-way ANOVA Sidak, n = 28 Con with 1-3 visits each, 15 IUGR with 1-3 visits each). Middle: Bayley Motor assessment scores across the first two years of life in Con and FGR infants (2-way ANOVA Sidak, n = 28 Con with 1-3 visits each, 15 IUGR with 1-3 visits each). Right: Bayley Language assessment scores across the first two years of life in Con and FGR infants (2-way ANOVA Sidak, n = 28 Con with 1-3 visits each, 15 IUGR with 1-3 visits each). **i,** Spearman correlation matrix of Shannon diversity index at week 3 and Staphylococcus genus % reads at week 2 against Bayley cognitive, motor, and language composite scores from 6 months (Con n = 10, FGR = 7; Con n = 10, FGR = 8, from left to right), 12 months (Con n = 8, FGR = 3; Con n = 7, FGR = 2, from left to right), and 18-24 months (Con n = 8, FGR = 4; Con n = 7, FGR = 4, from left to right). Displayed values are R^2^. Mean +/- SEM, *p < 0.05, **p < 0.01, ***p < 0.001, ****p < 0.0001, n.s. not significant.

Microorganisms from the maternal microbiota seed the infant gut microbiota at birth, serving as initial colonizers that inform infant metabolic and immune development^29^. We find in the maternal protein restriction mouse model for FGR that alterations in the maternal microbiota precede the development of neurobehavioral abnormalities in adult offspring. To gain insight into whether similar relationships are seen in a related human condition, we examined associations between the early life microbiome and neurocognitive outcomes in individuals from a cohort of preterm infants with or without FGR (**Supplementary Table 6-7**). Compared to 37 non-FGR preterm controls, 16 FGR preterm infants exhibited decreased alpha diversity of the fecal microbiota, with reduced Shannon diversity at week 3 of life, but no difference in subsequent weeks (**Fig. 3d, Supplementary Table 7**), suggesting delayed maturation of the gut microbiome. Taxonomic data did not show visible clustering of FGR vs non-FGR samples by principal coordinates analysis (**Fig. 3e**), indicating no widespread alterations in beta diversity. However, FGR samples exhibited significant alterations in select bacterial taxa, with increased *Staphylococcus* at 2 weeks of life (**Fig. 3f**). These alterations in microbiota were not attributable to statistically significant differences in infant sex, pregnancy induced hypertension, mode of delivery, day of life for first enteral feed, number of days on parenteral nutrition, antibiotics before or after delivery, early or late onset sepsis, or necrotizing enterocolitis (the latter showed a negative trending correlation with *Staphylococcus*) (**Fig. 3g, Supplementary Table 7**). When 28 non-FGR preterm infants were compared to 15 FGR preterm infants in a follow-up across 24 months corrected gestational age, the FGR subset scored significantly lower in the cognitive composite score on the Bayley Scales of Infant and Toddler Development III (BSID III), which is used to assess and diagnose developmental delays^61^ (**Fig. 3h**). Compared to non-FGR preterm controls, FGR preterm infants exhibited no statistical difference in language composite scores and a trending decrease in motor composite scores (**Fig. 3h**). In the subset of participants with both microbiota and BSID III outcome measures (15 non-FGR and 10 FGR), Shannon diversity at 3 weeks of life correlated positively with the language composite score at 12 months corrected gestational age and the cognitive composite score at 18-24 months corrected gestational age, while *Staphylococcus* at 2 weeks of life correlated negatively with cognitive composite score at 6, 12, and 18-24 months corrected gestational age, and trended towards a negative correlation with the motor composite score at 6 months corrected gestational age (**Fig. 3i**). *Staphylococcus*-dominated microbiota in preterm infants has previously been correlated to developmental delays and poor health outcomes^62^. Overall, these results reveal associations between human FGR, reduced alpha diversity of the early life gut microbiota, and reduced scores in infant neurodevelopmental measures, particularly in the cognitive domain. Despite the heterogenous nature and limited sample size of this human FGR cohort, and the difference in sampling timepoint, these findings mirror our observations from the maternal protein restriction mouse model for FGR.

### Depletion of the maternal gut microbiome modifies effects of dietary protein restriction on functional signatures in the fetal brain

The maternal gut microbiome guides normal fetal neurodevelopment and modifies effects of maternal environmental exposures, such as immune activation, antidepressant treatment, and prenatal stress on the fetal brain^24,25,58,63^. To examine influences of the maternal gut microbiome on fetal neurodevelopmental responses to maternal protein undernutrition, we examined functional signatures in the fetal brain after depletion of the maternal microbiome in PR-fed dams. Dams were treated with a cocktail of broad-spectrum antibiotics (ABX) by oral gavage twice daily for one week before breeding, and subsequently maintained on ABX in water throughout gestation^27,64^ (**Fig. 4a**). RNA sequencing revealed 1564 genes that were differentially expressed in E18.5 fetal brains from ABX-treated dams fed PR compared to conventional PR controls (specific pathogen free, SPF PR) (**Fig. 4b, Supplementary Table 2**), suggesting widespread fetal brain responses to depletion of the maternal microbiome. Of these, only 160 (10.2%) of the genes differentially expressed by maternal ABX treatment were similarly dysregulated by maternal PR relative to CD controls (**Fig. 4b**). By gene ontology analysis, fetal brain genes that were upregulated by ABX and PR mapped to pathways related to central nervous system development, neuron migration, synapse organization, and cellular response to amino acid starvation, whereas downregulated genes mapped to pathways including cerebral cortex and hippocampal development, as well as neuron migration (**Fig. 4c**). The minor overlap between the ABX and PR sets of differentially expressed genes suggests that the maternal microbiome has widespread influence on fetal brain gene expression that *modifies* responses to maternal protein restriction, but is unlikely to *mediate* adverse effects of maternal protein restriction.

**Figure 4:**
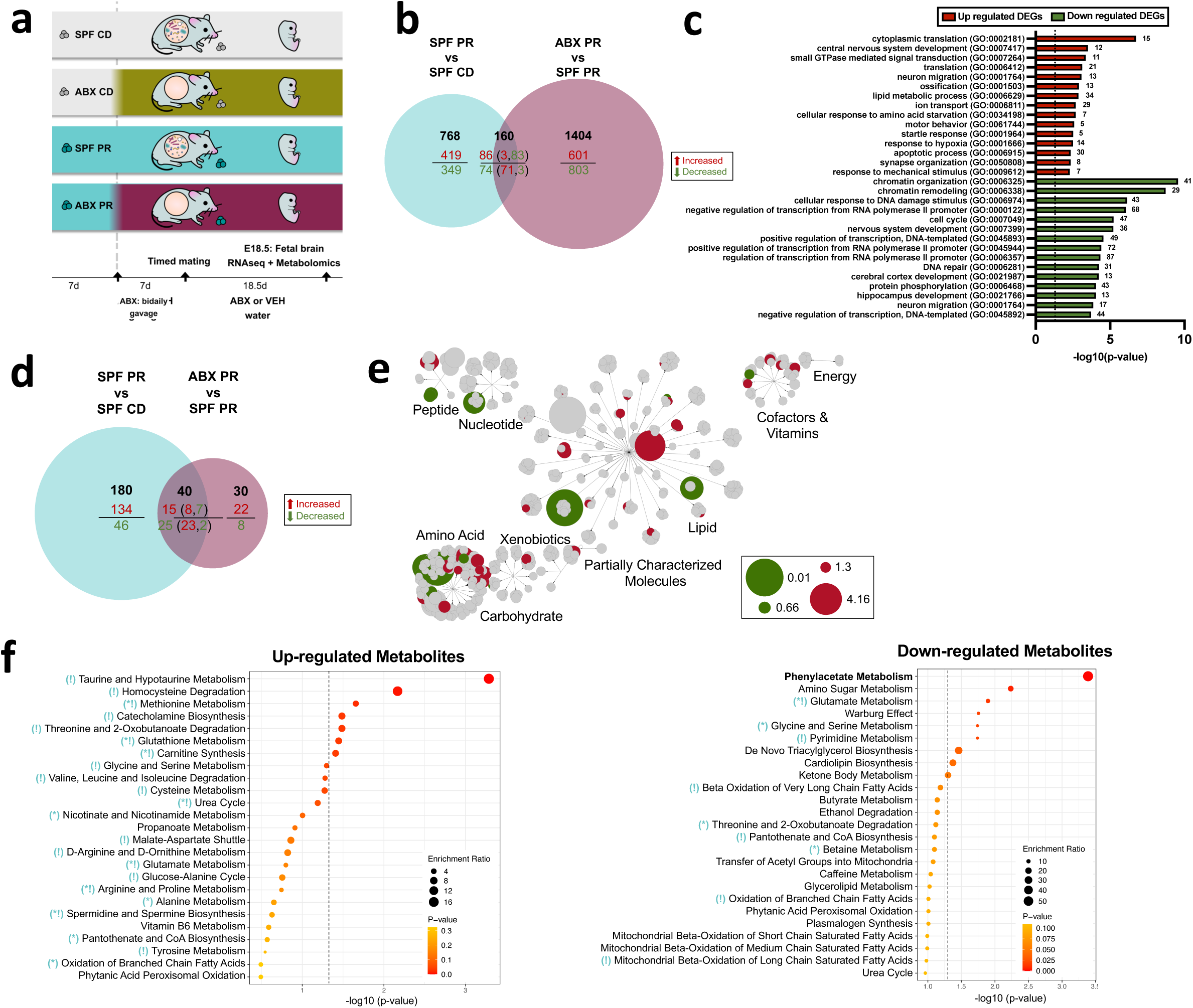
Depletion of the maternal gut microbiome induces distinct and additive fetal brain transcriptomic and metabolomic responses to maternal protein restriction. **a,** Timeline of ABX treatment prior to and throughout gestation. **b,** Venn diagram of DEGs (p < 0.05) in the fetal brains from SPF PR compared to SPF CD dams and in the fetal brains from ABX PR compared to SPF PR dams (n = 5 litters per group, 2 brains pooled per litter). Red = up-regulated, green = down-regulated. For the overlapping genes from both comparisons (center of the venn), red = up-regulated and green = down-regulated in SPF PR vs. SPF CD and in parentheses the colored numbers match the directionality of the metabolite in ABX PR vs SPF PR. **c,** Biological Process gene ontology (GO) of upregulated DEGs (p < 0.05, red) and downregulated DEGs (p < 0.05, green) of fetal brain transcripts from ABX PR dams and SPF PR dams (n = 5 litters per group, 2 brains pooled per litter). **d,** Venn diagram of differential metabolites (p < 0.05) in the fetal brains from SPF PR compared to SPF CD dams and in the fetal brains from ABX PR compared to SPF PR dams (n = 5 litters per group, 1.5-2 brains pooled per litter). Red = up-regulated, green = down-regulated. For the overlapping metabolites from both comparisons (center of the venn), red = up-regulated and green = down-regulated in SPF PR vs. SPF CD and in parentheses the colored numbers match the directionality of the metabolite in ABX PR vs SPF PR. **e,** Metabolon network map showing positive and negative fold change. **f,** Top enriched Small Molecule Pathway Database pathways for significantly up-regulated metabolites (p < 0.05, left) and down-regulated metabolites (p < 0.05, right) from fetal brains of ABX PR dams compared to SPF PR dams (bold pathways have q < 0.05; teal symbols relate to analogous enriched pathways in SPF PR vs SPF CD fetal brain [see Figure 2e]; * = enriched pathway in same direction; ! = enriched pathway in opposite direction).

In addition to modifying the fetal brain transcriptome, depleting the maternal microbiome in protein undernourished dams altered fetal brain metabolomic profiles. E18.5 brains from fetuses of antibiotic-treated PR dams exhibited significant alterations in 70 metabolites relative to SPF PR controls (**Fig. 4d**, **Supplementary Table 3**). Of these, 40 metabolites overlapped with those altered by maternal PR relative to CD controls, reflecting 18.2% of metabolites altered by PR relative to CD (**Fig 4d**). Metabolites altered by ABX were primarily amino acids (47.1%) and lipids (28.6%) (**Fig. 4e**), with significantly reduced metabolites mapping to 8 pathways, including phenylacetate metabolism (**Fig. 4f**). Maternal ABX-induced reductions in fetal brain metabolites related to glutamate metabolism, and glycine and serine metabolism (**Fig. 4f**), were similarly seen with maternal PR relative to CD (**Fig. 2e**), suggesting an exacerbating effect of maternal microbiome deficiency on these PR-induced alterations in the fetal brain. This finding is in line with work suggesting a role for the gut microbiome in supplying the host with essential amino acids^65^. Metabolites that were significantly elevated by depletion of the maternal microbiome mapped to 7 pathways, including taurine and hypotaurine metabolism and homocysteine degradation (**Fig. 4f**). Taken together, these results suggest that the diversity of the maternal microbiome alters the fetal brain in ways that modify effects of maternal protein undernutrition on offspring neurodevelopment.

### Microbiome-based interventions during pregnancy differentially modify adverse effects of maternal protein restriction on offspring growth or behavior

Alterations in the maternal microbiome during pregnancy are increasingly associated with impaired fetal and postnatal development of the offspring^27,30,31,66^, raising the question of whether manipulating the maternal microbiome during pregnancy can change offspring health trajectories. We found that maternal protein restriction alters maternal health status, including reductions in diversity of the maternal microbiota, which interacts with fetal development to yield abnormal behavior in adult offspring. Following the observation that ABX microbial depletion modifies PR-indued fetal brain signatures, we aimed to assess causal effects of the maternal microbiome during pregnancy on offspring developmental responses to maternal protein restriction. To this end, we investigated growth and behavior of offspring reared from ABX-treated and PR-fed dams, fed CD at birth, and cross-fostered to untreated SPF dams fed PR during pregnancy (**Fig. 5a**). This experimental design isolates the effect of maternal microbial depletion and PR to the pregnancy period, and evaluates their influences on offspring developmental programming. There was no significant effect of maternal microbial depletion on PR-induced reductions in fetal size, maternal gestational weight loss, or maternal diet consumption (**Extended Data Fig. 4a-c**). However, there was a partial attenuation of PR-induced elevation of maternal serum corticosterone levels at E18.5 (**Extended Data Fig. 4d**). This corresponded with a trending decrease in litter size (**Extended Data Fig. 4e**), despite overall increases in rates of litter survival at weaning (**Extended Data Fig. 4f**). Offspring of microbiota-depleted dams fed PR during pregnancy exhibited gradual sub-significant reductions in postnatal weight over the first two weeks of life (**Extended Data Fig. 4g**), with statistically significant decreases in weight by adulthood despite being fed CD since birth (**Extended Data Fig. 4h**). Maternal microbiome depletion and protein restriction during pregnancy yielded adult offspring with anxiety-like behavior that was not statistically different from that seen in control offspring from SPF and PR-fed dams (**Extended Data Fig. 5a** and 6a). However, adult offspring of ABX PR dams exhibited exacerbated learning deficits, with increased latency to target zone during the acquisition phase of the Barnes maze assay relative to cognitively impaired control offspring of SPF dams fed PR (**Fig. 5b**). In other parameters of the Barnes maze test, offspring of microbiome-depleted and PR-fed dams exhibited deficiencies in performance that were not statistically significantly different from those seen in control offspring from PR-fed SPF dams (**Extended Data Fig. 5b-d**). When analyzing data for male and female offspring separately, male offspring were especially affected by maternal microbiome depletion, with more pronounced exacerbation of cognitive behavior, which did not reach statistical significance (**Extended Data Fig. 6b-e**). These results indicate that further reducing the diversity of the maternal microbiome, as shaped by protein restriction during pregnancy, yields offspring developmental programs that lead to deficient postnatal weight gain and exacerbated cognitive impairment during adulthood. These findings further suggest that interventions to improve the diversity and function of the maternal microbiome may aid in preventing the adverse effects of maternal protein undernutrition on offspring growth and behavior.

**Figure 5:**
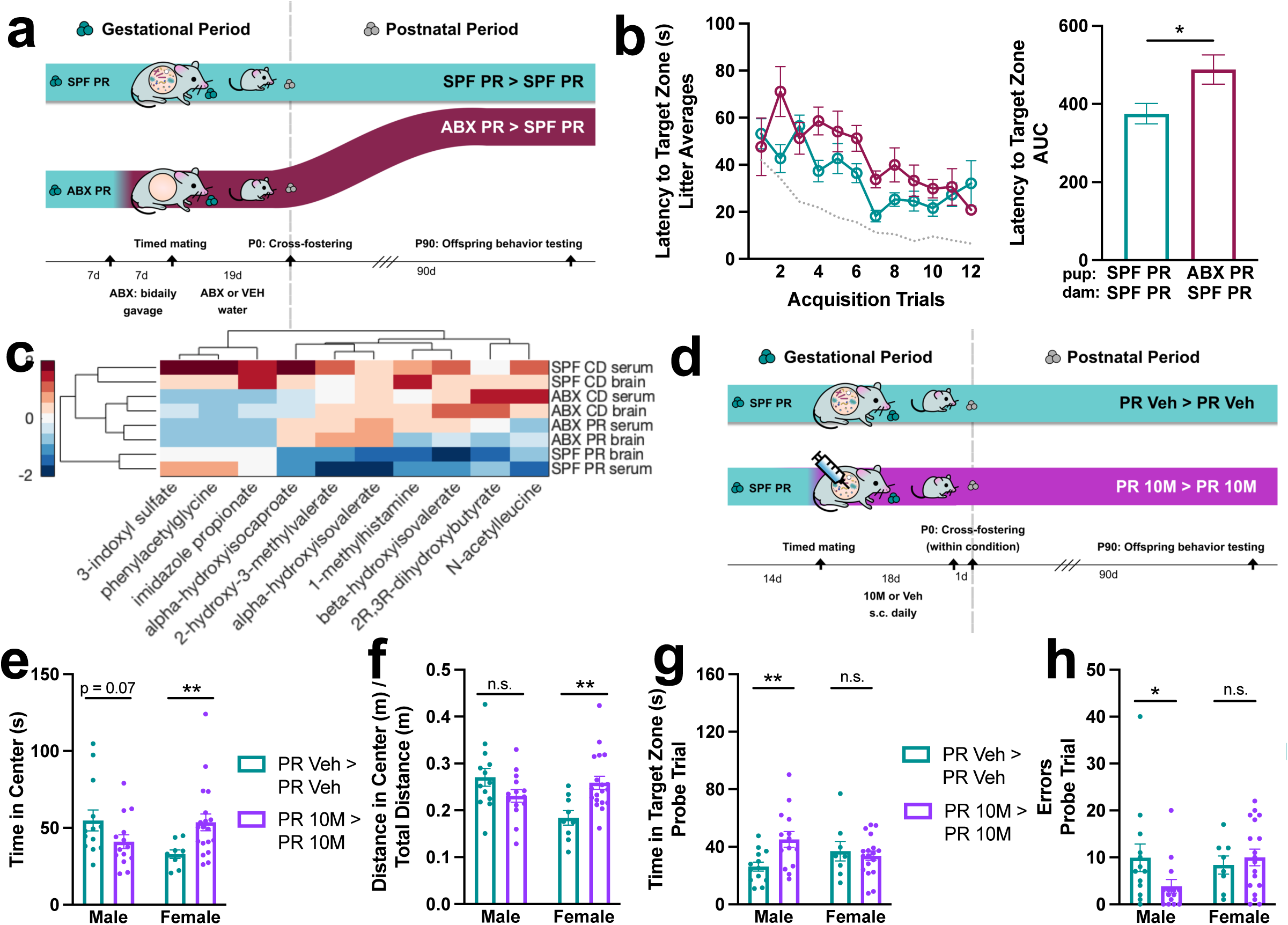
Maternal microbiome-informed interventions differentially modify risk for neurobehavioral deficits induced by maternal protein restriction during pregnancy. **a,** Graphic of cross-fostering paradigm for ABX experiments. **b,** Left: Latency to target zone in Barnes maze, all offspring averaged within each litter (n = 5 litters per group). Dotted line indicates average value for SPF CD litters. Right: AUC of latency to target zone (unpaired Welch’s t-test). Top row refers to pup condition, bottom row refers to dam condition. **c,** Heatmap of metabolites chosen for 10M supplementation, hierarchical clustering around 0, SD = 1 (n = 6 litters for each group/tissue type, 2 fetal brains or 1 dam serum pooled per litter). **d,** Graphic of cross-fostering paradigm for 10M experiments. **e,** Time in center in open field test, male and female offspring (Mann Whitney test for each sex, n = 13, 14, 9, 19, from left to right). **f,** Distance in center in open field test, controlled by total distance traveled, male and female offspring (Mann Whitney test for each sex, n = 13, 14, 9, 19, from left to right). **g,** Time in target zone in Barnes maze probe trial, male and female offspring (unpaired Welch’s t-test for each sex, n = 13, 14, 8, 18, from left to right). **h,** Errors in Barnes maze probe trial, male and female offspring (Mann-Whitney test for each sex, n = 13, 14, 8, 18, from left to right). Mean +/- SEM, *p < 0.05, **p < 0.01, ***p < 0.001, ****p < 0.0001, n.s. not significant.

Short chain fatty acids (SCFAs) produced by bacterial fermentation of complex carbohydrates promote normal gastrointestinal and immune function^67,68^, and their supplementation is reported to counter adverse effects of high fat or low fiber diets on behavioral development^55,56^. To explore potential microbiome-based interventions for preventing adverse effects of maternal protein undernutrition on offspring growth and behavior, we first tested supplementation with SCFAs, which we previously utilized to prevent placental insufficiencies induced by maternal protein restriction^64^. PR-fed dams were supplemented with a cocktail of SCFAs (acetate, butyrate, and propionate), or a sodium-matched vehicle control, in water throughout gestation^64^ (**Extended Data Fig. 7a**). There were no significant effects of maternal SCFA supplementation during pregnancy on maternal PR-induced reductions in fetal and maternal weight and elevations in maternal serum corticosterone, despite significant effects of SCFA supplementation on increasing maternal dietary intake (**Extended Data Fig. 7b-e**). Maternal SCFA supplementation had no statistically significant effects on litter size at birth, but modestly increased rates of litter survival postnatally, and promoted a gradual increase in pup weight over the first two weeks of life, which was reversed in adulthood (**Extended Data Fig. 7f-i**). Consistent with our previous observations (**Fig. 1**), adult offspring of vehicle-treated and gestational PR-fed dams exhibited anxiety-like behavior in the open field and cognitive impairment in the Barnes maze despite rearing since birth on CD (**Extended Data Fig. 8a-e**). There were no statistically significant effects of maternal SCFA supplementation on the behavioral abnormalities induced by maternal PR during pregnancy (**Extended Data Fig. 8a-e, 9a-e**). These results indicate that while maternal SCFA supplementation during pregnancy may effectively prevent adverse effects of maternal protein restriction on placental development^64^ and promote early postnatal growth (**Extended Data Fig. 7h**), it fails to avert diet-induced neurodevelopmental programming of behavioral abnormalities in adult offspring. This aligns with human studies, wherein children subject to early life protein undernutrition remain at risk for mood and neurocognitive disorders despite subsequent nutritional rehabilitation and adequate postnatal growth^13–16^.

The maternal microbiome regulates numerous metabolites in the maternal blood, fetal blood, and fetal brain, a subset of which aid in guiding normal fetal neurodevelopment^23,27,64^. We further explored the possibility that microbial metabolites aside from SCFAs could be used to attenuate adverse effects of maternal protein undernutrition on offspring development. To do so, we identified 10 microbially-modulated metabolites (10M: 3-indoxul sulfate, phenylacetylglycine, imidazole propionate, alpha-hydroxyisocaproate, 2-hydroxy-3-methylvalerate, alpha-hydroxyisovalerate, 1-methylhistamine, beta-hydroxyisovalerate, 2R, 3R-dihydroxybutyrate, and N-acetylleucine) that were reduced in the fetal brain by maternal PR, similarly reduced by maternal PR in maternal serum, and additionally altered by maternal ABX treatment, indicating dependence on the maternal microbiome (**Fig. 5c**). Pregnant dams were fed PR and treated with 10M, or vehicle control, throughout gestation via a daily subcutaneous injection (**Fig. 5d**). There were no statistically significant effects of maternal 10M supplementation on PR-induced reductions in fetal weight, maternal weight, and maternal food consumption (**Extended Data Fig. 10a-c**). However, maternal 10M supplementation significantly attenuated PR-induced increases in maternal serum corticosterone and drastically increased offspring survival (**Extended Data Fig. 10d**, f), with no effect on litter size at birth, pup weight, or adult weight (**Extended Data Fig. 10e**, g**-h**). Adult offspring of vehicle-treated dams fed PR exhibited anxiety-like behavior in the open field and cognitive behavioral impairment in the Barnes maze (**Extended Data Fig. 11a-e**), which reproduced effects of maternal protein undernutrition that were seen in initial PR experiments (**Fig. 1**) and in vehicle control groups for SCFA treatment (**Extended Data Fig. 8**). Adult offspring of 10M-supplemented and PR-fed dams displayed sexually dimorphic behavioral responses relative to control offspring of vehicle-treated and PR-fed dams (**Fig. 5e-h, Extended Data Fig. 12a-b**). In particular, female offspring of 10M-supplemented dams exhibited increased time and distance in the center during open field test without any motor effects (**Fig. 5e-f, Extended Data Fig. 11a**)), suggesting reduced anxiety-like behavior which aligns with the female-bias in anxiety-like behavior seen in response to maternal PR (**Extended Data Fig. 2b**). In contrast, male offspring of 10M-supplemented dams exhibited increased time in target zone and decreased errors during the probe phase of Barnes maze (**Fig. 5g-h**), suggesting better 24-hour recall without an effect on learning during the acquisition phase (**Extended Data Fig. 11b-e, 12a-b**). These results indicate that maternal 10M supplementation during pregnancy partially attenuates adverse effects of maternal protein undernutrition on abnormal cognitive and anxiety-related behavior in adult offspring in a sex-specific manner. These sex- and domain-specific improvements are seen in the absence of normalized fetal or postnatal growth, again providing evidence that amelioration of neurobehavioral trajectories can occur independently of restoration of physical growth.

## Discussion

Findings from this study identify the maternal microbiome as a modifier of adverse neurological outcomes in offspring of protein undernourished dams. Maternal protein restriction reduces diversity of the maternal microbiome and elicits widespread alterations in maternal-fetal metabolomic profiles and fetal brain gene expression, including subsets of maternal microbiome-dependent metabolites and genes. Further depleting the maternal microbiome of protein-restricted dams substantially alters transcriptomic and metabolomic profiles in fetal brains, where only a small fraction of metabolites and genes differentially regulated by maternal microbiome depletion overlap with those altered by protein restriction alone. Furthermore, depleting the maternal microbiome exacerbates cognitive, but not anxiety-like, behavioral impairments observed in adult offspring of protein-restricted dams. These results suggest that wholesale depletion of the maternal microbiome elicits select brain and behavioral changes in offspring of protein-restricted dams through mechanisms that are largely independent of those caused by maternal protein restriction alone. As such, reductions in microbial diversity likely modify, but do not directly mediate, adverse effects of maternal protein restriction on offspring neurodevelopment. These findings may have translational implications, as human preterm infants with FGR display early postnatal decreases in microbial diversity which correlate with reduced infant cognitive scores compared to non-FGR counterparts.

We find that supplementing protein-restricted dams with a cocktail of ten diet- and microbiome-dependent metabolites during pregnancy prevents anxiety-like behavior and cognitive impairment in their adult offspring. The metabolites were selected based on their significant decreases in both fetal brain and maternal serum of protein-restricted dams compared to standard protein-fed controls, and their further modulation by depletion of the maternal microbiome. It is unclear precisely how 10M is acting to protect offspring from behavioral impairments induced by maternal protein undernutrition. However, many of these metabolites have been previously shown to support neurological and behavioral functioning, in addition to being linked to the gut microbiome. Five of the ten metabolites (alpha-hydroxyisocaproate, 2-hydroxy-3-methylvalerate, alpha-hydroxyisovalerate, beta-hydroxyisovalerate, and N-acetylleucine) are dietary catabolites of the branched chain amino acids (BCAAs) leucine, isoleucine, and valine. The gut microbiota is capable of both producing and breaking down BCAAs^69^. BCAAs are also implicated in neural development and function following insult: alpha-hydroxyisocaproate, 2-hydroxy-3-methylvalerate, alpha-hydroxyisovalerate, and N-acetylleucine were increased in fetal brain following exposure to intrauterine inflammation^70^, and N-acetylleucine reduced cortical inflammation and apoptosis and increased memory in novel object recognition task after traumatic brain injury^71^. In addition to these five BCAA-related metabolites, 3-indoxyl sulfate, imidazole propionate, and phenylacetylglycine are well established metabolites of the gut microbiome produced by sulfonation of bacterially-derived indole, direct bacterial synthesis from histidine, and bacterial conversion of phenylalanine, respectively^72–74^. They have also been implicated in neurodevelopment: imidazole propionate promoted thalamocortical axonogenesis in fetal brains from offspring of antibiotic-depleted dams^27^, and all three have been reported to vary across postnatal development in mouse forebrain^75^. The final 10M metabolites, 1-methylhistamine and 2R,3R-dihydroxybutyrate (commonly referred to as 4-deoxyerythronic acid), have only correlational ties to the gut microbiome and brain development. The former, a major metabolite of histamine, has been correlated with microbes, primarily of the order *Clostridiales* and genus *Lactobacillus*^76^, while the latter, a metabolite of L-threonine^77^, was downregulated in the plasma of MDD patients after escitalopram treatment^78^. Specific microbial metabolites are increasingly linked to neurodevelopment and behavior. In addition to the examples discussed above, individual microbially-modulated metabolites have recently been reported to induce anxiety-like behaviors by preventing oligodendrocyte maturation and altering myelination^79^, and promote cognitive decline by precipitating microglial apoptosis^80^. However, more research is needed to identify neuromodulatory microbial metabolites, and further understand the diverse mechanisms by which they engage in direct or indirect communication with the brain during critical temporal windows to guide neurodevelopment and influence behavioral outcomes.

Although the exact mechanisms by which PR induces behavioral deficits, and by which 10M ameliorates them are unclear, transcriptomic and metabolomic profiling of fetal brains of PR-fed dams revealed some possibilities. For one, tryptophan and tryptophan-related metabolic pathways were significantly downregulated in maternal serum, but significantly upregulated in fetal brain in response to gestational PR. Additionally, the *Slc6a4* gene encoding the serotonin transporter and the *Htr3a* gene encoding a serotonin receptor are significantly decreased in PR fetal brains. Finally, a tryptophan catabolite, 3-indoxyl sulfate, was among the 10M group that normalized behavior in offspring. These findings suggest abnormal tryptophan and serotonin signaling pathways as candidate contributors to subsequent behavioral deficits in offspring. Indeed, tryptophan and serotonin were altered by protein restriction during gestation and lactation in rats^81^, serotonin regulates many neurodevelopmental processes, and altered levels during gestation have been linked to behavioral impairments, including anxiety and cognitive, in offspring^82^. In addition, disruptions to the hypothalamic-pituitary-adrenal axis could be at play, as we observe elevated corticosterone in maternal serum and in fetal brains of protein-restricted dams, the former of which is attenuated by 10M supplementation. Stress during critical periods can be neurotoxic, is known to interact with both maternal microbiome and fetal neurodevelopment^25,59^, and to precipitate anxiety- and cognitive-related behavioral deficits in offspring^83^. Finally, neuroimmune processes may be disrupted, as we observed downregulated T-cell differentiation in PR fetal brains by transcriptomic pathway analysis. Indeed, T-cell differentiation promoted by maternal gut bacteria has been reported to underlie behavioral deficits in offspring following maternal immune activation^24,84^. Although we interpret these changes to be due to low levels of protein in the PR diet, high-sucrose diets during gestation have also been shown to alter offspring brain and behavior^85,86^. While the PR diet has comparably lower sucrose levels than used in high-sucrose models (15% kcal coming from increased sucrose and cellulose in PR compared to CD, versus 25% kcal from sucrose alone in a high-sucrose model^85^), and we see minimal carbohydrate metabolites affected in PR maternal serum and fetal brains, sucrose is significantly higher in PR fetal brain compared to CD. We are therefore unable to rule out an effect of high sucrose on the reported results.

This study showed sexual dimorphism in behavioral responses to both PR alone, where female offspring showed a more severe anxiety-like phenotype, and to 10M intervention, where female offspring showed restoration of anxiety-like behavior, and male offspring showed restoration of memory deficits. These findings highlight the importance of examining both males and females, and including sex as a biological variable^87^. Sex biases have been well-documented in various neurodevelopmental conditions, including a male skew in autism spectrum disorder and attention deficit hyperactivity disorder^88^ and a female skew in anxiety and mood disorders^89^. Furthermore, findings from this study support the ability of sex to influence susceptibility to, or presentation of, prenatal programming from environmental stressors or maternal microbial shifts^59^. The biological basis for this interaction with sex is unclear, but theories have been raised to explain sexually dimorphic responses to prenatal perturbations, including slower maturation and increased intrauterine immunoreactive responses to male fetuses, differences in placental function by sex, and contributions of sex steroids and *SRY* programming^90,91^. These different presentations of behavioral domain by sex also raise the question of different biological underpinnings of anxiety-like versus cognitive behaviors. Projections from the basolateral amygdala to the central amygdala^92^ and ventral hippocampus^93^ underlie anxiety-related behaviors, while spatial learning and memory rely on hippocampal CA3-medial prefrontal cortex circuits^94^. Based on the sexually dimorphic and domain-specific behavioral responses of offspring treated with 10M during gestation, it may be the case that these interventions are acting in a sex- and circuit-specific manner to ameliorate select phenotypes of gestational protein restriction. There is more work to be done in understanding the role of microbes and microbial metabolites in supporting or disrupting neurodevelopment, and how biological sex interacts with and facilitates this process.

Although maternal supplementation with 10 diet- and microbiome-dependent metabolites prevented impairments in particular anxiety-like and cognitive behavioral parameters in adult offspring, not all behavioral deficits induced by maternal protein restriction were prevented, suggesting that they are mediated by additional, as yet undefined, microbiome-dependent or - independent mechanisms. Results from our cross-fostering experiments indicate that maternal protein restriction alters offspring behavior through mechanisms that require diet-induced programming of both fetal neurodevelopment and maternal physiology, as only offspring born to and fostered by dams on PR during gestation show behavioral deficits. One way in which maternal programming could translate is via maternal care behaviors, which are well-known to inform offspring neurodevelopment and behavior^50,51^. Although we do not see significant differences in maternal retrieval behavior, which is altered in other gestational stress models^95^, there may be changes in other domains of stress-sensitive maternal care that warrant further investigation, such as licking/grooming and nursing behaviors^95,96^. Additionally, milk volume or nutrient content may be persistently altered by gestational PR, and may therefore create inconsistencies in postnatal nutrition between cross-fostering groups to inform long-term behavioral trajectories. Indeed, protein content of milk and size of offspring was reduced in dams fed a low protein diet during gestation and lactation^97^, and sialylated milk oligosaccharides, found to be decreased in mothers of undernourished infants, were sufficient to promote growth in humanized mouse and pig models of early postnatal malnutrition^98^. The impact of malnutrition and other peri-gestational stressors on both maternal and infant health underscores the importance of further research on women’s health and investigation of lasting implications of the postpartum period.

Nutritional support, and even growth and gross physiological restoration, are often inadequate to prevent long-term microbial, neurological, and behavioral impairments caused by early malnutrition^5–7,12–16^. Indeed, results from this study show consistent decoupling of early growth trajectories with later behavioral impairments. Restricting protein undernutrition to the pregnancy period in mice induces anxiety-like and impaired cognitive behavior in adult offspring that are fed standard nutritive diet since birth and exhibit typical growth trajectories by body weight. Depletion of the maternal microbiome exacerbates, whereas maternal supplementation with select microbes or microbially modulated metabolites prevents, select behavioral deficits in adult offspring without influencing pre- and postnatal growth. In contrast, maternal supplementation with short-chain fatty acids promotes feto-placental^64^ and offspring growth, but does not effectively prevent neurobehavioral deficits in adult offspring of dams that were protein restricted during pregnancy. As malnutrition continues to be a prevalent and severe global health burden^23^, additional interventions beyond nutritional rehabilitation alone are needed to target adverse effects of malnutrition on child neurobehavioral development. Findings from this study reveal that the maternal microbiome modifies adverse neurobehavioral outcomes of gestational protein undernutrition, which are partially prevented by maternal supplementation with select microbial metabolites. These results suggest that microbiome-based interventions should be further evaluated for their ability to reduce malnutrition-associated neurobehavioral disorders, which are not adequately addressed with current nutrition-focused standards of care.

## Methods

### Mice

8-week old male and female C57Bl/6J mice were purchased from Jackson Laboratories and maintained on 12-h light-dark cycle in temperature- and humidity-controlled environment. All mice were kept under sterile conditions (autoclaved cages, bedding, water bottles, and water). All experiments were performed in accordance with the NIH Guide for the Care and Use of Laboratory Animals using protocols approved by the Institutional Animal Care and Use Committee at UCLA.

#### Protein restriction

Mice were first subjected to a microbiota normalization, where two scoops of bedding were removed from each cage, mixed together, and deposited back in each cage to normalize microbiota between all mice in each testing cohort. At least 5 days after normalization, male and female mice were given either 6% protein diet (Teklad Envigo; TD.90016) or 20% control diet (Teklad Envigo; TD.91352) *ad libitum* for at least 2 weeks. Following this habituation period, mice were paired and time-mated. At E0.5, determined by observation of copulation plug, pregnant dams were individually housed and continued to be fed the same diet through gestation. Weights and fecal samples were collected from each dam at E0.5 and 18.5, with a subset collected at E0.5, 7.5, 10.5, 14.5, 18.5.

#### Antibiotic treatment

For “ABX” treatment groups, SPF male and female mice were gavaged twice daily for one week with neomycin (100 mg/kg), metronidazole (100 mg/kg) and vancomycin (50 mg/kg), as previously described^27,64^. During this time, ampicillin (1 mg/mL) was provided *ad libitum* in drinking water. Mice were then time-mated as described above. Following gavage, mice were maintained on ampicillin (1 mg/mL), neomycin (1 mg/mL) and vancomycin (0.5 mg/mL) *ad libitum* in drinking water for the duration of the experiment.

#### Fetal tissue collections

At E18.5, dams were sacrificed by cervical dislocation. Fecal samples were extracted from the colon. Blood was collected by cardiac puncture into vacutainer SST tubes (Becton Dickinson) and allowed to clot for 1 hour at room temperature, before centrifuging at 1500xg, 4°C for 10 minutes. Serum supernatant was collected and stored at -20°C. The entire uterine horn, including all conceptuses, was removed and placed in ice cold 1x PBS. Dams were weighed once again to record post-transection weight. Each fetus was dissected from the amniotic sac and weighed. Brains were microdissected and either placed into RNAlater (Invitrogen) for subsequent RNA sequencing, or snap-frozen in liquid nitrogen for subsequent metabolomics analysis. All samples were stored at -80°C.

### Behavioral testing

For behavioral cohorts, dams were given shepherd shacks on E17.5 to reduce stress ahead of birth. At P0, dams and diet were weighed, pups were counted, and all diet groups were switched to CD. Complete litters were cross-fostered, as denoted in figures (**Fig. 1a**, **Fig. 5a,d, Extended Data Fig. 7a**). Subsequently, litters were checked daily for pup survival. Litters were denoted as not survived if <4 pups remained at weaning (as previously done^64^). Pups were weighed on P3, 5, 7, 9, 11, 13, and toe-tattooed P4-P6, for identification purposes. Pups were weaned at P24-P26, and separated into cages with same-sex littermates. Adult behaviors were run beginning when mice were ∼90 days old. All mice were habituated in the testing room 30 minutes prior to test start. Tests were run in order of least to most stressful, in the order described below, with at least 3 days separating each test. All equipment was cleaned with 70% ethanol and allowed to dry in between trials.

#### Righting reflex & maternal retrieval – SPF cohorts only

The righting reflex and maternal retrieval tests were utilized to examine early neurological development^99^ and maternal care-giving and responsiveness^53^, respectively. Pups were subjected to righting reflex and maternal retrieval tests on P3, 5, 9, 11. All pups from each litter were placed one-by-one on their backs and the time it took for them to right themselves (with all four paws on the ground) was recorded, capped at 30 seconds. Pups were then placed back into their home cage all at once, in the opposite corner of the nest. The latency to first pup retrieval and full litter retrieval by the dam was measured, capped at 5 minutes.

#### Ultrasonic vocalizations – SPF cohorts only

Ultrasonic vocalizations in pups were analyzed as a response to separation from the litter ^100^. Pups were tested for USVs at P7 and P13. Four pups were randomly chosen (when possible, 2 males and 2 females) for USV testing, and the same pups were used at both timepoints. Pups were habituated in the recording chambers for 5 minutes, and then recorded for 5 minutes. Number and duration of calls, in addition to inter-call latency, were analyzed. Recording and analysis were performed using SASLab Pro (Avisoft Bioacoustics).

#### Open field test – all cohorts

Open field test was employed to assess locomotion and thigmotaxis as a proxy for anxiety-like behavior^35,36^. Mice were placed into open field arenas (27.5 cm^2^) for 10 minutes under bright light. Videos were recorded using an overhead camera, and the first 5 minutes of each test were analyzed using AnyMaze 7.1 for time spent in center zone, distance traveled in center zone/total distance traveled, total distance traveled, and mean speed.

#### Barnes maze – all cohorts

Barnes maze was employed to assess spatial learning and memory^38^. The Barnes maze apparatus has an elevated round surface (90 cm diameter) with 20 holes evenly spaced around the perimeter. One hole was denoted as the “escape” hole, and an escape box was attached below the hole. The position of the escape hole was kept consistent within mice, but counterbalanced between mice. Spatial cues (black and white geometric shapes printed on 9x11in paper) were placed around the maze, and kept consistent throughout training and testing. On day 1 of the testing protocol, each mouse was allowed to freely explore the Barnes maze apparatus for 5 minutes with no white noise under mild light, with no escape box present. Immediately following, each mouse was placed under a wire cup in the center of the apparatus, with white noise and bright light for 1 minute. Finally, each mouse was gently guided to the escape hole and into the escape box, and white noise and bright light were removed, for 1 minute. On days 2-5, each mouse was subject to three training trials, with an inter-trial latency of 20-45 minutes. The training trials consisted of 15s habituation, where mice were placed under a wire cup in the center of the apparatus with bright light and white noise. The cup was then lifted, and mice were given 180s to escape into the escape hole, thereby ending the test. If the mouse did not escape by 180s, they were gently guided into the escape hole.

At the end of each trial, mice were left for 1 minute in the escape hole, with no white noise. On day 6, a probe test was run to assess long-term memory. Each mouse was subject to one probe trial, consisting of a 15s habituation, and then a 180s test without the escape box. Videos were recorded using an overhead camera and analyzed in AnyMaze 7.1. For the training trials, latency to find the escape hole and latency to enter the escape box were analyzed. For the probe trial, time spent in the escape hole vicinity, and errors, defined as non-escape hole vicinities a mouse entered *before* entering the escape hole vicinity, were analyzed.

### Human Infants

This prospective cohort study was conducted at the University of California Los Angeles (UCLA) neonatal intensive care units (UCLA Ronald Reagan Hospital/UCLA Mattel Children’s Hospital, Los Angeles, CA and UCLA Santa Monica Hospital, Santa Monica, CA). The UCLA Institutional Review Board granted approval for this study (IRB #15 00718). Verbal informed consent was obtained from parents/legal guardians. Inclusion criteria included infants with a birth weight <2 kilograms (kg), <14 days of age, who were predicted to require parenteral nutrition for ≥2 weeks, and follow-up at the UCLA High Risk Infant Clinic (HRIF). The exclusion criteria included infants who were unlikely to survive and infants with congenital anomalies and congenital infections. Infants missing prenatal medical records with which to diagnose FGR were also excluded.

Demographic and clinical data was collected from the electronic medical chart. FGR was determined as previously described^101^ by the obstetric team who documented fetal growth deceleration on repeated prenatal ultrasounds. Full feeds were defined as 100 mL/kg/d of enteral nutrition or *ad libitum* feeding, whichever occurred first. Early onset sepsis was defined as a positive blood culture before 72 hours of age and antibiotics for ≥5 days. Late onset sepsis was defined as a positive blood culture after 72 hours of age and antibiotics for ≥5 days. Necrotizing enterocolitis was defined by Bell’s stage II or III. Neurodevelopment was assessed using composite cognitive, language, and motor scores from the Bayley Scales of Infant Development (BSID) III^61^, which was conducted by doctorally trained clinicians who work in the HRIF. These clinicians have established inter-rater reliability for the BSIS-III examination, which is monitored for recertification research projects and clinical assessments..

### 16S rRNA gene sequencing

Fecal samples were collected from CD and PR dams throughout gestation at the following timepoints: E0.5, E7.5, E10.5, E14.5, E18.5 and stored at -80°C until processing. The same dams were collected from at each timepoint. Bacterial genomic DNA was isolated according to manufacturer instructions using the DNeasy PowerSoil Kit (Qiagen). Additionally, fecal samples were collected from preterm infants with and without FGR during their hospital stay. Specimens were collected shortly after study enrollment, then weekly while on parenteral nutrition, and for four weeks after parenteral nutrition was discontinued. Samples were collected from infant diapers using sterile collection kits and stored at -80°C until processing. Because of intestinal dysmotility, preterm infants have delayed passage of meconium and do not stool frequently in the first couple weeks of life, resulting in missing samples disproportionately at early timepoints. Approximately 50mg per sample was aliquoted in liquid nitrogen, bead-beat in buffer RLT using Lysing Matrix E tubes (MP Biomedicals), and extracted using AllPrep DNA/RNA/Protein Mini kit (Qiagen), as previously described^102^.

For both mouse and human samples,16S rRNA gene sequencing was performed as previously described^27,64^. Briefly, sequencing libraries were generated according to methods adapted from Caporaso et al. 2011^103^, amplifying the V4 regions of the 16S rRNA gene by PCR. The final PCR product was purified using the Qiaquick PCR purification kit (Qiagen). 250 ng of purified, PCR product from each individually barcoded sample were pooled and sequenced by Laragen, Inc. using the Illumina MiSeq platform and 2 x 250bp reagent kit for paired-end sequencing. All analyses were performed using QIIME2 2023.7^104^, including DADA2^105^ for quality control, taxonomy assignment, alpha-rarefaction and beta-diversity analyses. 29,152 reads were analyzed per sample. Operational taxonomic units (OTUs) were assigned based on 99% sequence similarity compared to the SILVA 138 database. Beta-diversity principal coordinates analysis plots were generated with QIIME2 View, and other bacterial metrics, including correlation analyses with clinical metadata were performed and plotted in Prism 9.0 (Graphpad).

### Corticosterone ELISAs

Total corticosterone levels were determined by a corticosterone ELISA assay (Enzo Life Sciences). The assay was run per manufacturer’s instructions, following the small volume protocol. All experimental samples were run in triplicate; blanks and controls were run in duplicate. Mean optical density was read on a Synergy H1 plate reader (Agilent BioTek), and corticosterone concentrations were determined in ug/mL based on a standard curve.

### Fetal brain RNA sequencing

Whole fetal brains were randomly chosen from each litter, dissected, and immediately placed into 400uL RNA*later* (ThermoFisher Scientific) and stored at -80°C. RNA was extracted and cDNA synthesized using the PureLink RNA Mini Kit (Invitrogen) and qScript cDNA synthesis kit (Quantabio), respectively. Equal amounts of RNA were pooled from 2 brains per litter, resulting in 1522ng RNA per sample, except for one which had 508ng. An RNA integrity number (RIN) of at least 8.7 was confirmed for each sample using the 4200 Tapestation system (Agilent). RNA was prepared as previously described^64^ using the TruSeq RNA Library Prep kit and Lexogen QuantSeq 3’ forward-sequencing was performed using the Illumina HiSeq 4000 platform by the UCLA Neuroscience Genomics Core. Sequences were quality filtered, trimmed, and mapped using FastQC v0.11.5, bbduk v35.92, and RSeQC v2.6.4. Reads were aligned to UCSC Genome Browser assembly ID: mm10 using STAR v2.5.2a, indexed using samtools v1.3, and aligned using HTSeq-count v0.6.0. Differential expression analysis was conducted using DESeq2 v1.24.041. Heatmaps were generated using the pheatmap v1.0.12 package for R. DAVID 2021^106,107^ was used to conduct GO term enrichment analysis of differentially expressed genes with non-adjusted p-value < 0.05.

### Metabolomics

Whole fetal brains were randomly chosen from each litter, dissected, and immediately snap-frozen in liquid nitrogen and stored at -80°C. 1.5-2 brains were pooled for subsequent analysis. Untargeted metabolomics was run on fetal brain and maternal serum by Metabolon Inc. as previously described^27,64^. Briefly, samples were prepared using the automated MicroLab STAR system (Hamilton Company) and analyzed on GC/MS, LC/MS and LC/MS/MS platforms. Organic aqueous solvents were used to perform serial extractions for protein fractions, concentrated using a TurboVap system (Zymark) and vacuum dried. For LC/MS and LC-MS/MS, samples were reconstituted in acidic or basic LC-compatible solvents containing > 11 injection 25 standards and run on a Waters ACQUITY UPLC and Thermo-Finnigan LTQ mass spectrometer, with a linear ion-trap frontend and a Fourier transform ion cyclotron resonance mass spectrometer back-end. For GC/MS, samples were derivatized under dried nitrogen using bistrimethyl-silyl-trifluoroacetamide and analyzed on a Thermo-Finnigan Trace DSQ fast-scanning single-quadrupole mass spectrometer using electron impact ionization. Chemical entities were identified 30 by comparison to metabolomic library entries of purified standards. Following log transformation and imputation with minimum observed values for each compound, data were analyzed using two-way ANOVA to test for group effects. P- and q-values were calculated based on two-way ANOVA contrasts. Principal components analysis was used to visualize variance distribution with the ggplot2 R package^108^. The Metaboanalyst 5.0^109^ platform’s metabolite set enrichment analysis (MSEA) was performed on whole fetal brain and maternal serum metabolites that were statistically significantly altered (non-adjusted p-value < 0.05) between SPF PR and SPF CD treatment groups and ABX PR and SPF PR treatment groups. Metabolites sets were analyzed for metabolite pathway enrichment using the Small Molecule Pathway Database (SMPDB).

### In vivo short-chain fatty acid supplementation

All mice were habituated on PR diet, and bred as described above. Once copulation plugs were observed, SCFA were supplemented *ad libitum* in drinking water from E0.5-E18.5 (prenatal cohort) or E19.5 (behavior cohort), as previously described^64^. SCFA water cocktail contained 25mM sodium propionate, 40mM sodium butyrate, and 67.5mM sodium acetate. SCFA water was changed at least every 4 days. Sodium-matched controls were given water supplemented with 7.745g/L sodium chloride. All supplemented water was sterile-filtered before administration. SCFA and Na-matched litters were then used for E18.5 tissue collections, or for postnatal cross-fostering and behavioral testing, as described above.

### In vivo metabolite supplementation

#### Metabolite selection for in vivo supplementation

Metabolites were first filtered for significant differences between SPF CD and SPF PR. Those that were decreased in SPF PR compared to SPF CD were further checked for microbiome modulation, as denoted by significant change in any direction in ABX PR relative to SPF PR, or in ABX CD relative to SPF CD. This set of filters was applied to metabolomic results from both fetal brain and maternal serum. Metabolites that met these criteria from *both* tissue compartments were used for supplementation.

#### Metabolite supplementation

All mice were habituated on PR diet, and bred as described above. Once copulation plugs were observed, sterile filtered metabolite mixture (10M) was injected subcutaneously once per day from E0.5-E17.5. Control dams were injected with vehicle. Physiological concentrations were determined as previously described^27^ based on literature^72,110–118^, estimated blood volume, and on the relative differences in metabolite fold change between CD and PR dams (**Supplementary Tables 3-4**). The 10M injections contained 0.76ug imidazole propionate, 7.3ug alpha-hydroxyisocaproate, 7.24ug alpha-hydroxyisovalerate, 0.0094uL beta-hydroxyisovalerate, 11.68ug N-acetylleucine, 2.16ug 2-hydroxy-3-methylvalerate, 6.3ug phenylacetylglycine, 29.16ug 3-indoxyl sulfate, 0.04ug 1-methylhistamine, and 6.66ug 2R,3R-dihydroxybutyrate in 200uL 1X sterile PBS. Vehicle injections contained 8.66ug potassium chloride, 0.012nL hydrochloric acid, and 1.7ug sodium hydroxide in 200uL 1X sterile PBS. 10M and vehicle litters were then used for E18.5 tissue collections, or for postnatal cross-fostering and behavioral testing, as described above.

### Statistical Methods and Sample Sizes

Statistical analyses were performed using Prism 9.0 (Graphpad). Assays with two groups were assessed for normality and subsequently analyzed by either an unpaired two-tailed t-test with Welch’s correction, or by a Mann-Whitney test. Tests on three or more groups were assessed by a one-way ANOVA, and when there were two factors, with a two-way ANOVA. Data over time were assessed with a repeated measures ANOVA or mixed measures analysis. Tukey or Sidak post-hoc tests were used, based on Prism recommendations. All prenatal weight measures and postnatal behavioral measures were run using litter as a biological replicate, with all fetuses or offspring averaged within each litter. For prenatal weights and maternal metrics, at least 9 litters per group were analyzed. For behavioral experiments, 4-7 litters per group were tested. To assess sex differences in the behavioral results, and in adult weights, individual male and female offspring were treated as biological replicates. For metabolomics, 1.5-2 fetal brains were randomly chosen and pooled per litter, from 6 litters per group. For RNAseq, 2 fetal brains were randomly chosen and pooled per litter, from 5 litters per group. Whole litters were excluded from all analyses if they had fewer than 4 fetuses or pups (as done previously^64^). For prenatal measures, no viable fetuses within any litters were excluded from reported analyses. For behavioral measures, individual offspring were occasionally excluded from all tests due to health reasons (i.e. developing malocclusion) or from a specific test or trial due to a test-related event (i.e. escaping the testing chamber). These occasions were rare, and not associated with a particular group or condition. All data are plotted as mean +/- SEM. Significant differences are denoted as follows: *p < 0.05, **p < 0.01, ***p < 0.001, ****p < 0.0001, n.s. not significant.

### Data Availability

Transcriptomic data that support the findings are available in Supplementary Table 2 and have been deposited to the Sequence Read Archive (SRA) repository with accession number PRJNA1074327. Untargeted metabolomic data that support the findings are available in Supplementary Tables 3 and 4 and have been deposited to Mendeley Data (DOI: 10.17632/2pdwjm85tb.1. 16S rRNA gene sequencing data that support the findings are available in Supplementary Tables 5 and 6 and have been deposited to the SRA repository with accession number PRJNA1074327. Human clinical metadata that support the findings are available in Supplementary Table 7. Other data available upon reasonable request.

## Acknowledgements

We thank members of the Hsiao laboratory for their guidance and review of the manuscript; members of the UCLA Goodman Luskin Microbiome Center Gnotobiotics Core Facility for technical support; members of the laboratory of Dr. Grace Aldrovandi for their assistance with human 16S sequencing; Dr. Lindsay Lueptow and Irina Zhuravka of the UCLA Behavioral Testing Core for critical training on mouse behavioral testing; Joe DeYoung and Helen Ibsen of the UCLA Neuroscience Genomics Core for their assistance with RNA sequencing; Dr. Stephanie White for allowing usage of USV recording equipment; Dr. Lars Bode for helpful insights regarding postnatal milk; and Drs. Carlos Portera-Cailliau, Sherin Devaskar, Bennet Novitch, and Bridget Callaghan for helpful feedback on the project.

## Contributions

E.J.C.O. led the study, performed the experiments, and analyzed the data. G.R.L assisted with analyzing datasets from 16S rRNA gene sequencing, transcriptomics, and metabolomics. G.N.P. assisted with timed matings and provided key technical guidance on establishing the PR model.

E.O. and K.B.Y. assisted with 16S analysis and gnotobiotic interpretations. J.M., A.C., and E.G. assisted with behavioral and ELISA assays. H.E.V. assisted with timed matings and provided key technical guidance. J.P. assisted with mouse experiments and provided technical guidance on gnotobiotic work. A.C. and K.L.C led the human FGR study. E.J.C.O. and E.Y.H. designed the study and wrote the manuscript. All authors discussed the results and commented on the manuscript.

## Ethics Declarations

The authors declare no competing interests.

---

**Extended Data Figure 1:**
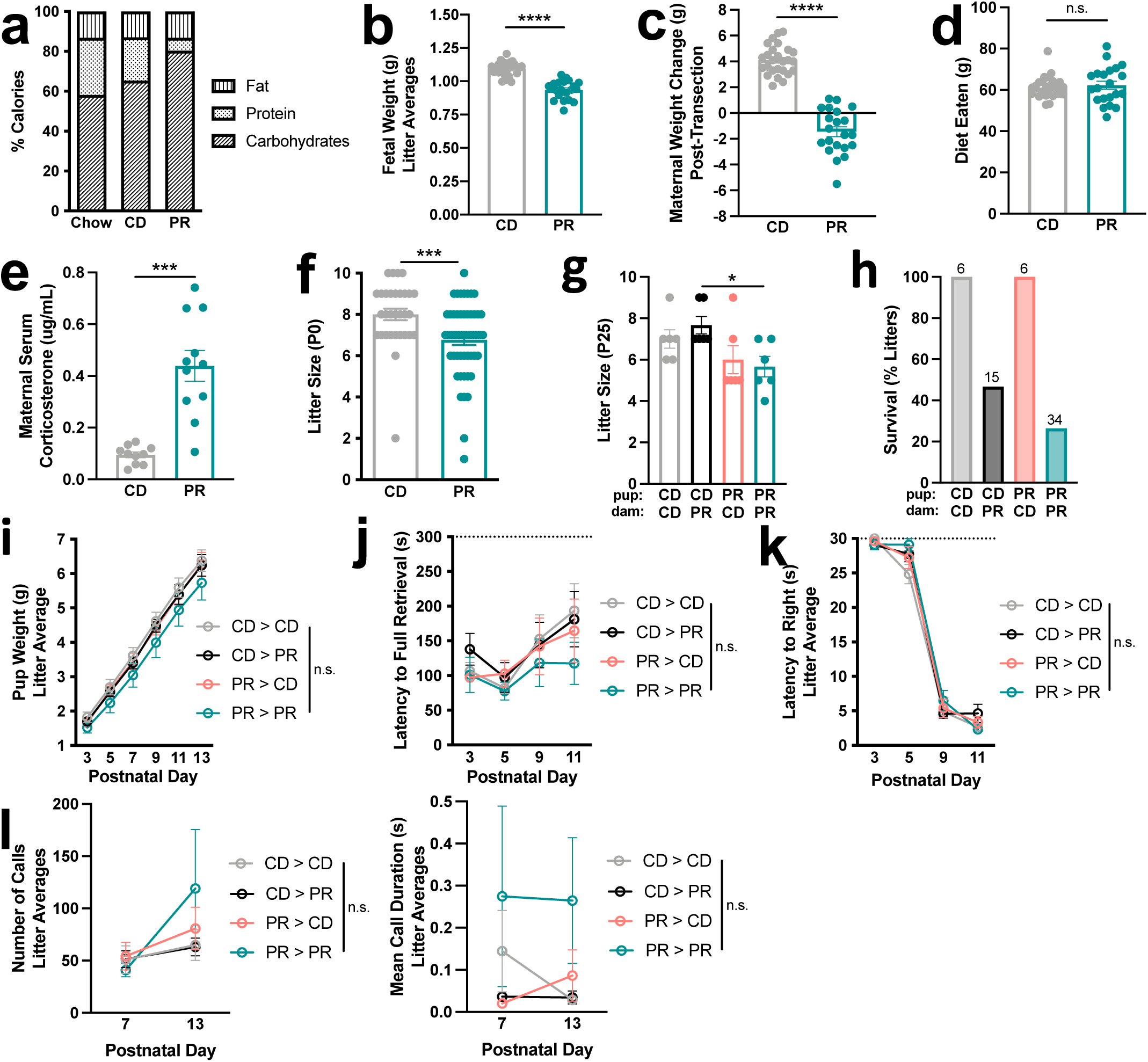
Maternal protein restriction induces fetal growth restriction and maternal stress, and reduces postnatal survival, but does not influence early postnatal growth or maternal behavior. **a,** Macronutrient breakdown of CD and PR diets compared to standard lab chow. **b,** Fetal weight at E18.5 from SPF CD and SPF PR dams, all fetuses averaged within each litter (unpaired Welch’s t-test, n = 28, 21, from left to right). **c,** Maternal weight change, from E0.5 to E18.5, in SPF CD and SPF PR dams (unpaired Welch’s t-test, n = 26, 21, from left to right). **d,** Maternal diet eaten, from E0.5 to E18.5 in SPF CD and SPF PR dams (Mann Whitney test, n = 26, 21, from left to right). **e,** Corticosterone measured in serum in SPF CD, SPF PR dams at E18.5 (unpaired Welch’s t-test, n = 10, 11, from left to right). **f,** Litter size at P0, from SPF CD and SPF PR dams (Mann Whitney test, n = 31, 50, from left to right). **g,** Litter size, pups per litters, measured at weaning, from cross-fostered groups SPF CD pups > CD dams, SPF CD pups > PR dams, SPF PR pups > CD dams, SPF PR pups > PR dams (two-way ANOVA with Sidak, n = 6 litters per group). Top row refers to pup condition, bottom row refers to dam condition. **h,** Litter survival (percentage of total litters), from SPF CD pups > CD dams, SPF CD pups > PR dams, SPF PR pups > CD dams, SPF PR pups > PR dams (n = 6, 15, 6, 34, from left to right). Top row refers to pup condition, bottom row refers to dam condition. **i,** Pup weights, all offspring averaged within each litter (two-way repeated measures ANOVA with Tukey, n = 6 litters per group). **j,** Latency to right in pup righting reflex test, all offspring averaged within each litter (two-way repeated measures ANOVA with Tukey, n = 6 litters per group). **k,** Latency to retrieve in maternal retrieval test (two-way repeated measures ANOVA with Tukey, n = 6 dams per group). **l,** Left: Number of calls, ultrasonic vocalizations, 4 offspring averaged within each litter (two-way repeated measures ANOVA with Tukey, n = 6 litters per group). Right: Mean duration of calls, ultrasonic vocalizations, 4 offspring averaged within each litter (two-way repeated measures ANOVA with Tukey, n = 6 litters per group). Mean +/- SEM, *p < 0.05, **p < 0.01, ***p < 0.001, ****p < 0.0001, n.s. not significant.

**Extended Data Figure 2:**
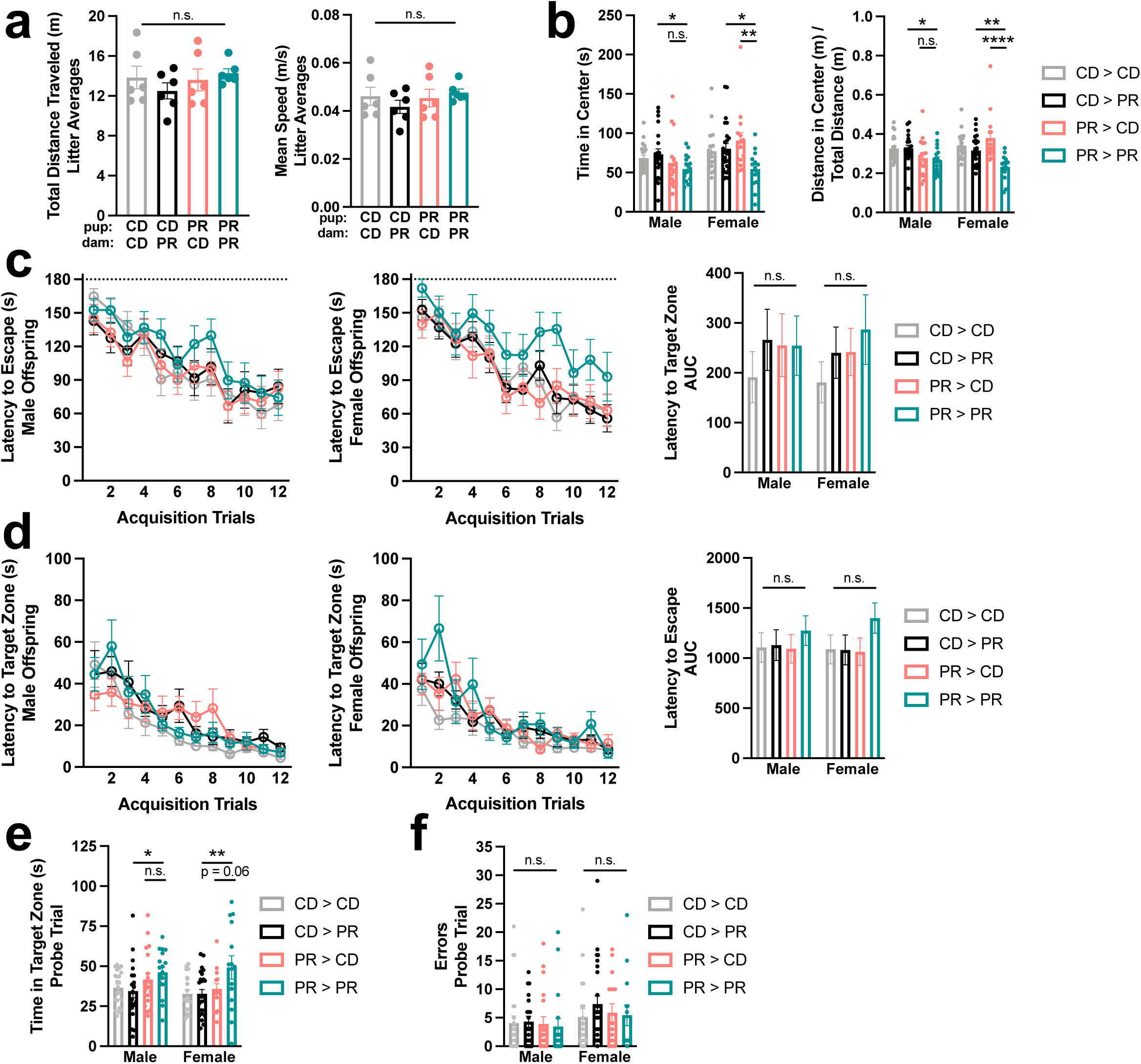
Adult female offspring exhibit behavioral differences based on both gestational and rearing-associated protein restriction, whereas male offspring are based on gestational alone. **a,** Left: Total distance traveled in open field test, all offspring averaged within each litter (two-way ANOVA with Sidak, n = 6 litters per group). Right: Mean speed in open field test, all offspring averaged within each litter (two-way ANOVA with Sidak, n = 6 litters per group). Top row refers to pup condition, bottom row refers to dam condition. **b,** Left: Time in center in open field test, male and female offspring (two-way ANOVA with Sidak for each sex, n = 22, 20, 18, 18, 19, 25, 15, 14, from left to right). Right: Distance in center in open field test, controlled by total distance traveled, male and female offspring (two-way ANOVA with Sidak for each sex, n = 22, 20, 18, 18, 19, 25, 15, 14, from left to right). **c,** Left: Latency to escape in Barnes maze acquisition phase, male offspring (n = 22, 20, 19, 18). Middle: Latency to escape in Barnes maze acquisition phase, female offspring (n = 19, 25, 15, 14). Right: AUC of latency to escape (two-way ANOVA with Sidak for each sex). **d,** Left: Latency to target zone in Barnes maze acquisition phase, male offspring (n = 22, 20, 19, 18). Middle: Latency to target zone in Barnes maze acquisition phase, female offspring (n = 19, 25, 15, 14). Right: AUC of latency to target zone (two-way ANOVA with Sidak for each sex). **e,** Time in target zone in Barnes maze probe trial, male and female offspring (two-way ANOVA with Sidak for each sex, n = 22, 20, 19, 18, 19, 25, 15, 14, from left to right). **f,** Errors made in Barnes maze probe trial, male and female offspring (two-way ANOVA with Sidak for each sex, n = 22, 20, 19, 18, 19, 25, 15, 14, from left to right). Mean +/- SEM, *p < 0.05, **p < 0.01, ***p < 0.001, ****p < 0.0001, n.s. not significant.

**Extended Data Figure 3:**
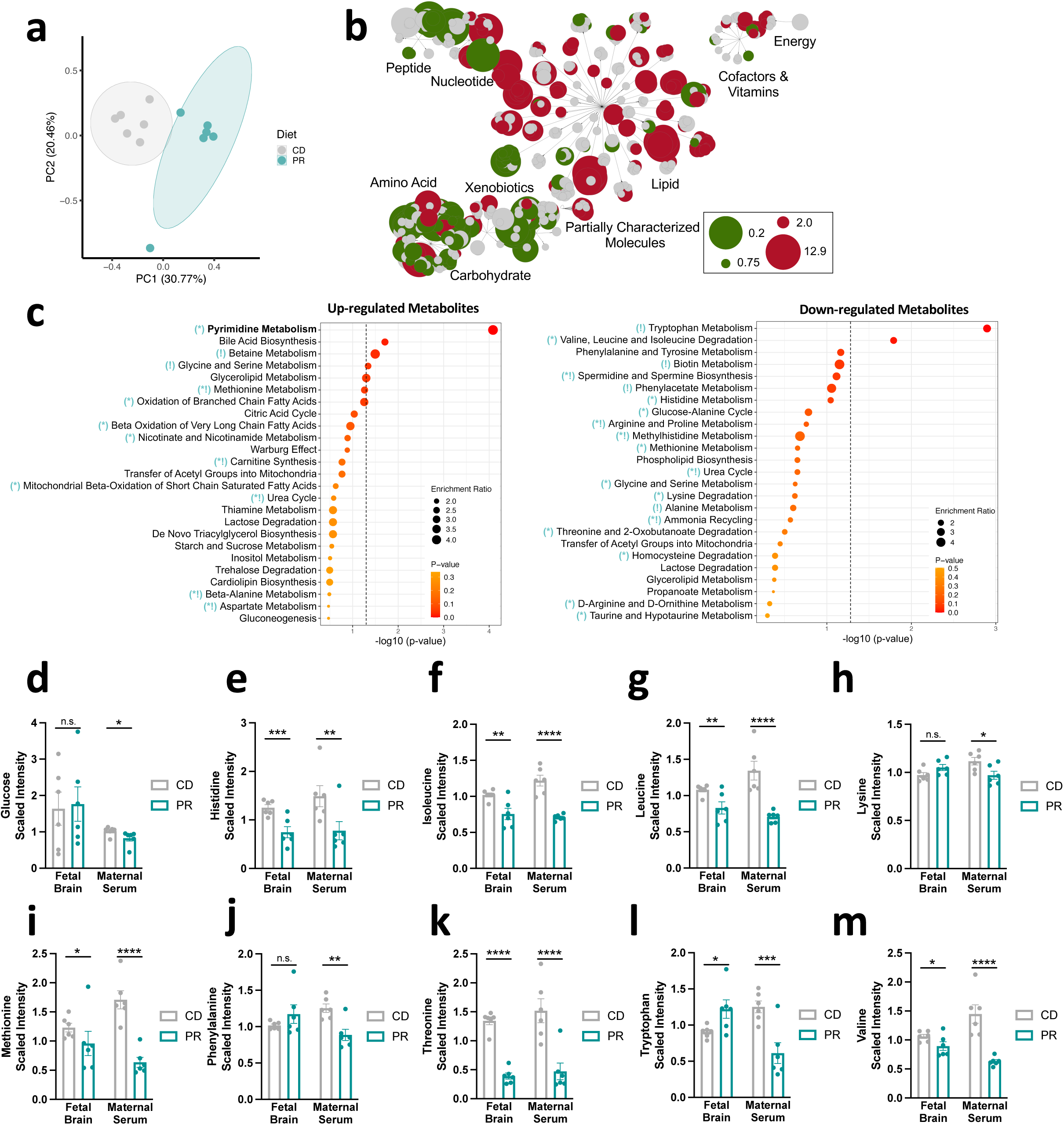
Maternal protein restriction alters metabolomic profiles in maternal serum, and induces nutrient brain sparing in fetal brains at late gestation. **a,** Untargeted metabolomics PCA comparing serum from SPF CD and SPF PR dams (n = 6 dams per group). **b,** Metabolon network map showing positive and negative fold change. **c,** Top enriched Small Molecule Pathway Database pathways for significantly upregulated metabolites (p < 0.05, left) and downregulated metabolites (p < 0.05, right) for serum from SPF PR dams compared to SPF CD dams (bold pathways have q < 0.05; teal symbols relate to analogous enriched pathways in SPF PR vs SPF CD fetal brain [see Figure 2e]; * = enriched pathway in same direction; ! = enriched pathway in opposite direction). **d-m,** Untargeted metabolomics of essential amino acids and glucose from E18.5 fetal brains from SPF CD and SPF PR dams (n= 6 litters per group, 1.5-2 brains pooled per litter) and E18.5 serum from SPF CD and SPF PR dams (n = 6 dams per group). Mean +/- SEM, *p < 0.05, **p < 0.01, ***p < 0.001, ****p < 0.0001, n.s. not significant.

**Extended Data Figure 4:**
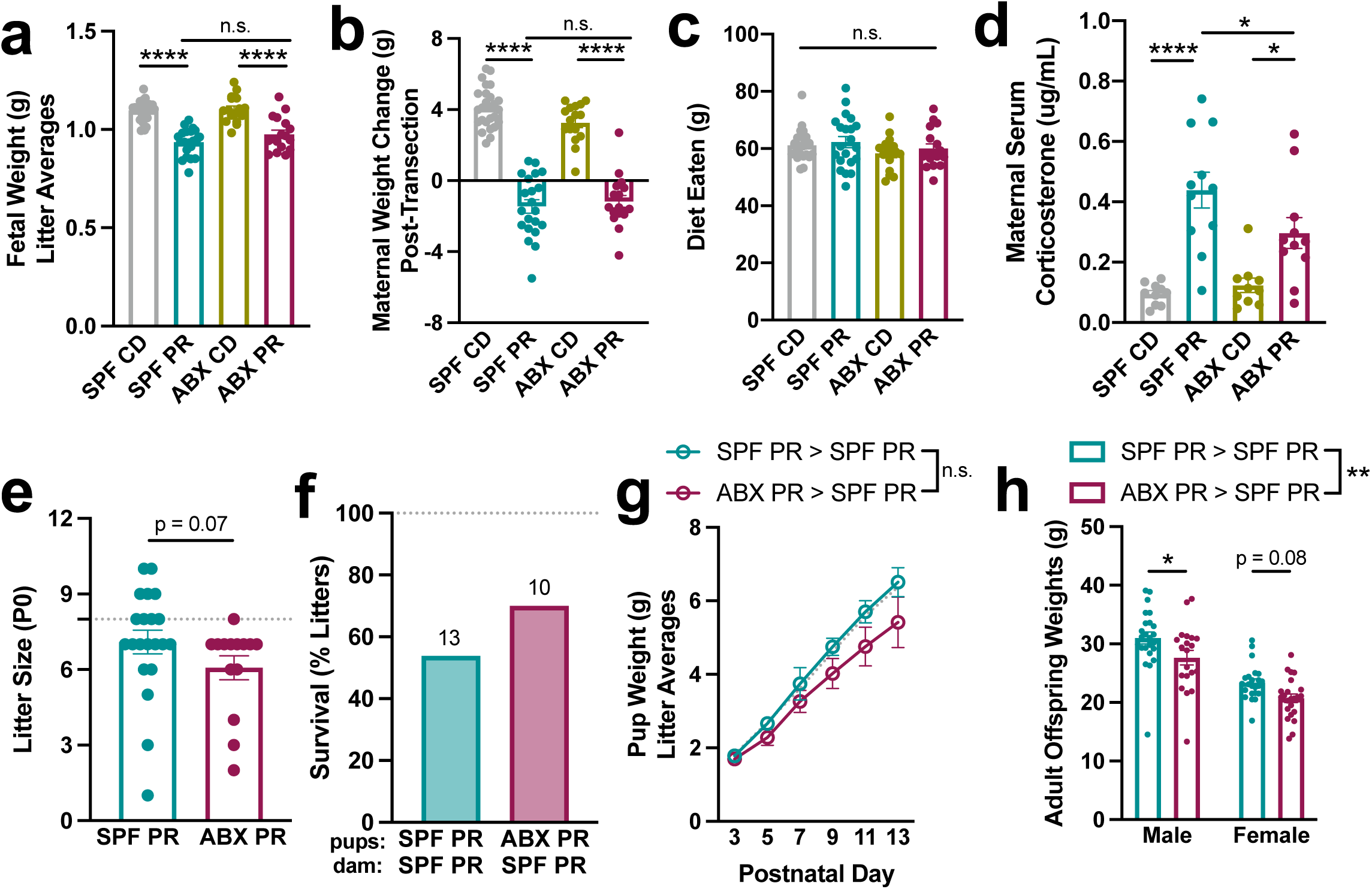
Maternal microbial depletion moderately impacts gross measures of maternal and offspring health. **a,** Fetal weight at E18.5 from SPF CD, SPF PR, ABX CD, ABX PR dams, all fetuses averaged within each litter (two-way ANOVA with Sidak, n = 28, 21, 16, 17, from left to right). SPF data same as in Extended Data Fig. 1b. **b,** Maternal weight change, from E0.5 to E18.5 post-transection, in SPF CD, SPF PR, ABX CD, ABX PR dams (two-way ANOVA with Sidak, n = 26, 21, 16, 17, from left to right). SPF data same as in Extended Data Fig. 1c. **c,** Diet eaten, from E0.5 to E18.5, in SPF CD, SPF PR, ABX CD, ABX PR dams (two-way ANOVA with Sidak, n = 26, 21, 16, 17, from left to right). SPF data same as in Extended Data Fig. 1d. **d,** Corticosterone measured in serum in SPF CD, SPF PR, ABX CD, ABX PR dams at E18.5 (two-way ANOVA with Sidak, n = 10, 11, 10, 11, from left to right). SPF data same as in Extended Data Fig. 1e. **e,** Litter size, pups per litters, measured at P0, from SPF PR and ABX PR dams (Mann-Whitney test, n = 21, 14, from left to right). Dotted line indicates average value for SPF CD litters. **f,** Litter survival (percentage of total litters), from SPF PR pups > SPF PR dams and ABX PR pups > SPF PR dams (n = 13, 10, from left to right). Dotted line indicates average value for SPF CD litters. Top row refers to pup condition, bottom row refers to dam condition. **g,** Pup weights, all offspring averaged within each litter (two-way mixed effects analysis with Sidak, n = 7 litters per group). Dotted line indicates average value for SPF CD litters. **h,** Adult weights, male and female offspring (two-way ANOVA with Sidak, n = 24, 20, 23, 23, from left to right). Mean +/- SEM, *p < 0.05, **p < 0.01, ***p < 0.001, ****p < 0.0001, n.s. not significant.

**Extended Data Figure 5:**
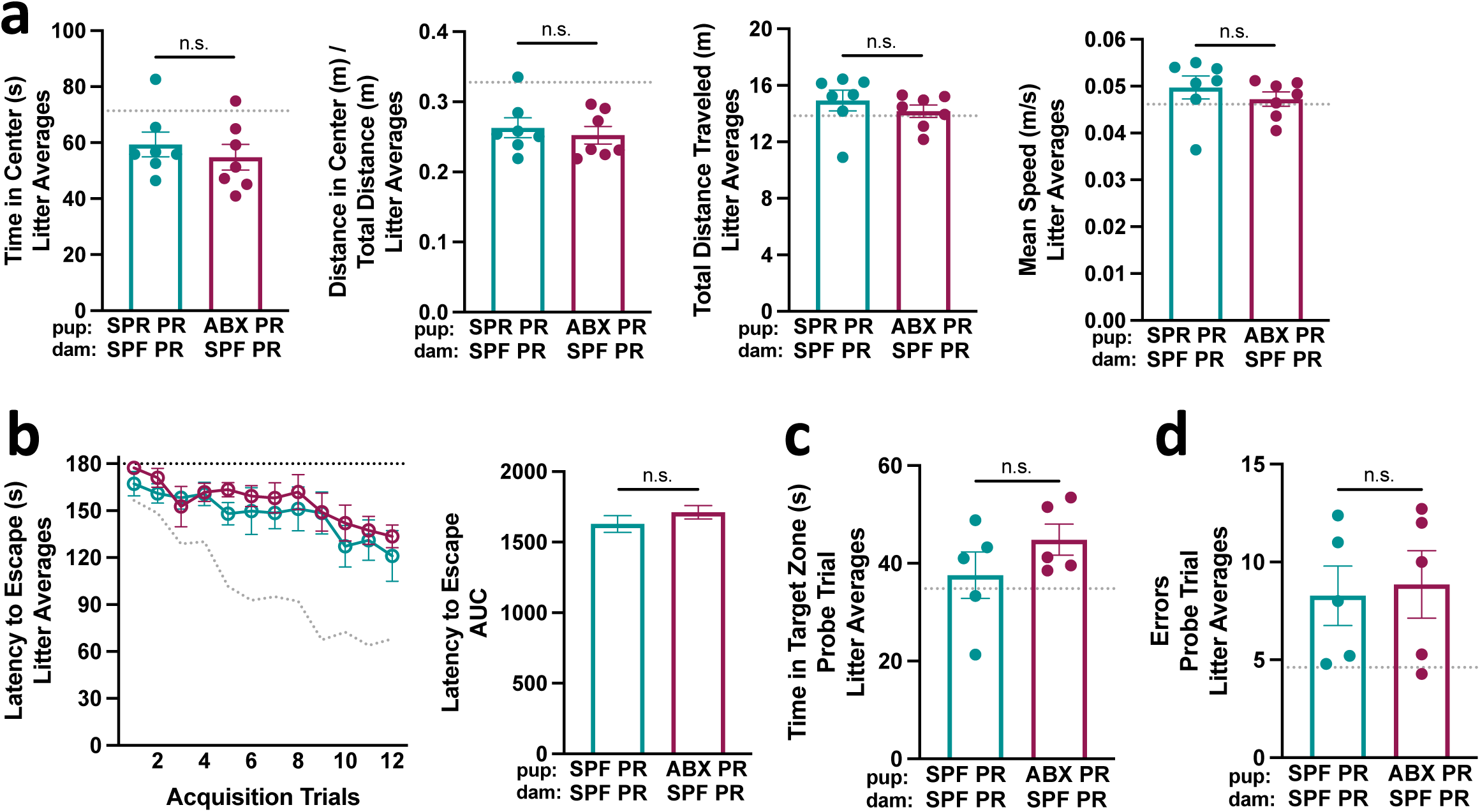
Maternal microbiome depletion does not influence anxiety-like, locomotor, or cognitive behavioral measures in adult offspring exposed to gestational protein restriction. **a,** Left: Time in center in open field test, all offspring averaged within each litter (Mann Whitney test, n = 7 litters per group). Top row refers to pup condition, bottom row refers to dam condition. Dotted line indicates average value for SPF CD litters. Left-middle: Distance in center in open field test, controlled by total distance traveled, all offspring averaged within each litter (unpaired Welch’s t-test, n = 7 litters per group). Top row refers to pup condition, bottom row refers to dam condition. Dotted line indicates average value for SPF CD litters. Right-middle: Total distance traveled in open field test, all offspring averaged within each litter (Mann Whitney test, n = 7 litters per group). Top row refers to pup condition, bottom row refers to dam condition. Dotted line indicates average value for SPF CD litters. Right: Mean speed in open field test, all offspring averaged within each litter (Mann Whitney test, n = 7 litters per group). Top row refers to pup condition, bottom row refers to dam condition. Dotted line indicates average value for SPF CD litters. **b,** Left: Latency to escape in Barnes maze, all offspring averaged within each litter (n = 5 litters per group). Dotted line indicates average value for SPF CD litters. Right: AUC of latency to escape (unpaired Welch’s t-test). Top row refers to pup condition, bottom row refers to dam condition. **c,** Time in target zone in Barnes maze probe phase, all offspring averaged within each litter (unpaired Welch’s t-test, n = 5 litters per group). Dotted line indicates average values for SPF CD litters. **e,** Errors made in Barnes maze probe phase, all offspring averaged within each litter (unpaired Welch’s t-test, n = 5 litters per group). Dotted line indicates average values for SPF CD litters. Mean +/- SEM, *p < 0.05, **p < 0.01, ***p < 0.001, ****p < 0.0001, n.s. not significant.

**Extended Data Figure 6:**
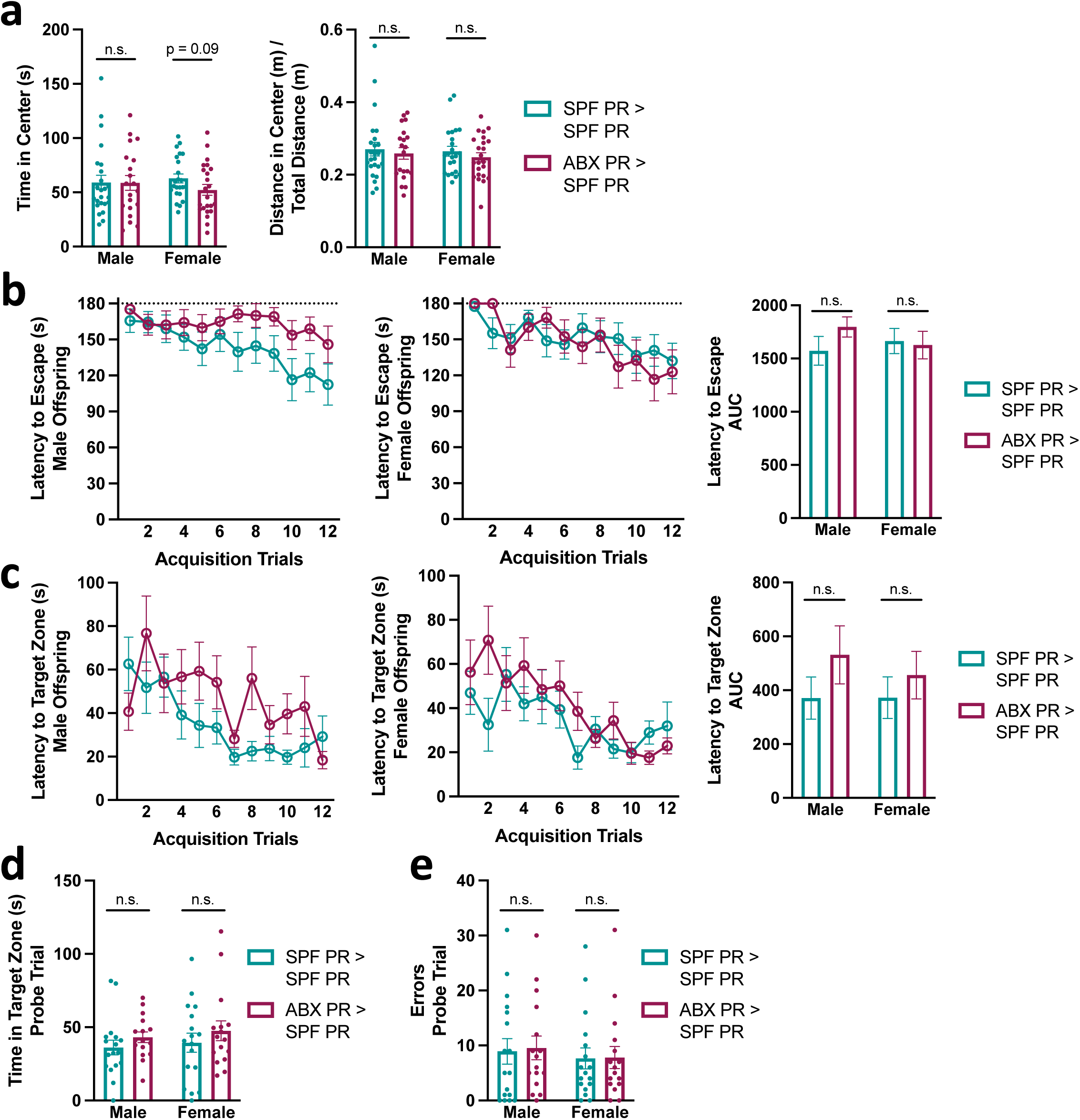
Adult offspring do not exhibit sexually dimorphic behavioral responses to gestational protein restriction and maternal microbiome depletion. **a,** Left: Time in center in open field test, male and female offspring (Mann Whitney test for each sex, n = 24, 20, 23, 23, from left to right). Right: Distance in center in open field test, controlled by total distance traveled, male and female offspring (Mann Whitney test for each sex, n = 24, 20, 23, 23, from left to right). **b,** Left: Latency to escape in Barnes maze acquisition phase, male offspring (n = 17, 16). Middle: Latency to escape in Barnes maze acquisition phase, female offspring (n = 17, 16). Right: AUC of latency to escape (unpaired Welch’s t-test for each sex). **c,** Left: Latency to target zone in Barnes maze acquisition phase, male offspring (n = 17, 16). Middle: Latency to target zone in Barnes maze acquisition phase, female offspring (n = 17, 16). Right: AUC of latency to target zone (unpaired Welch’s t-test for each sex). **d,** Time in target zone in Barnes maze probe trial, male and female offspring (Mann-Whitney test for each sex, n = 17, 17, 16, 16, from left to right). **e,** Errors made in Barnes maze probe trial, male and female offspring (Mann-Whitney test for each sex, n = 17, 17, 16, 16, from left to right). Mean +/- SEM, *p < 0.05, **p < 0.01, ***p < 0.001, ****p < 0.0001, n.s. not significant.

**Extended Data Figure 7:**
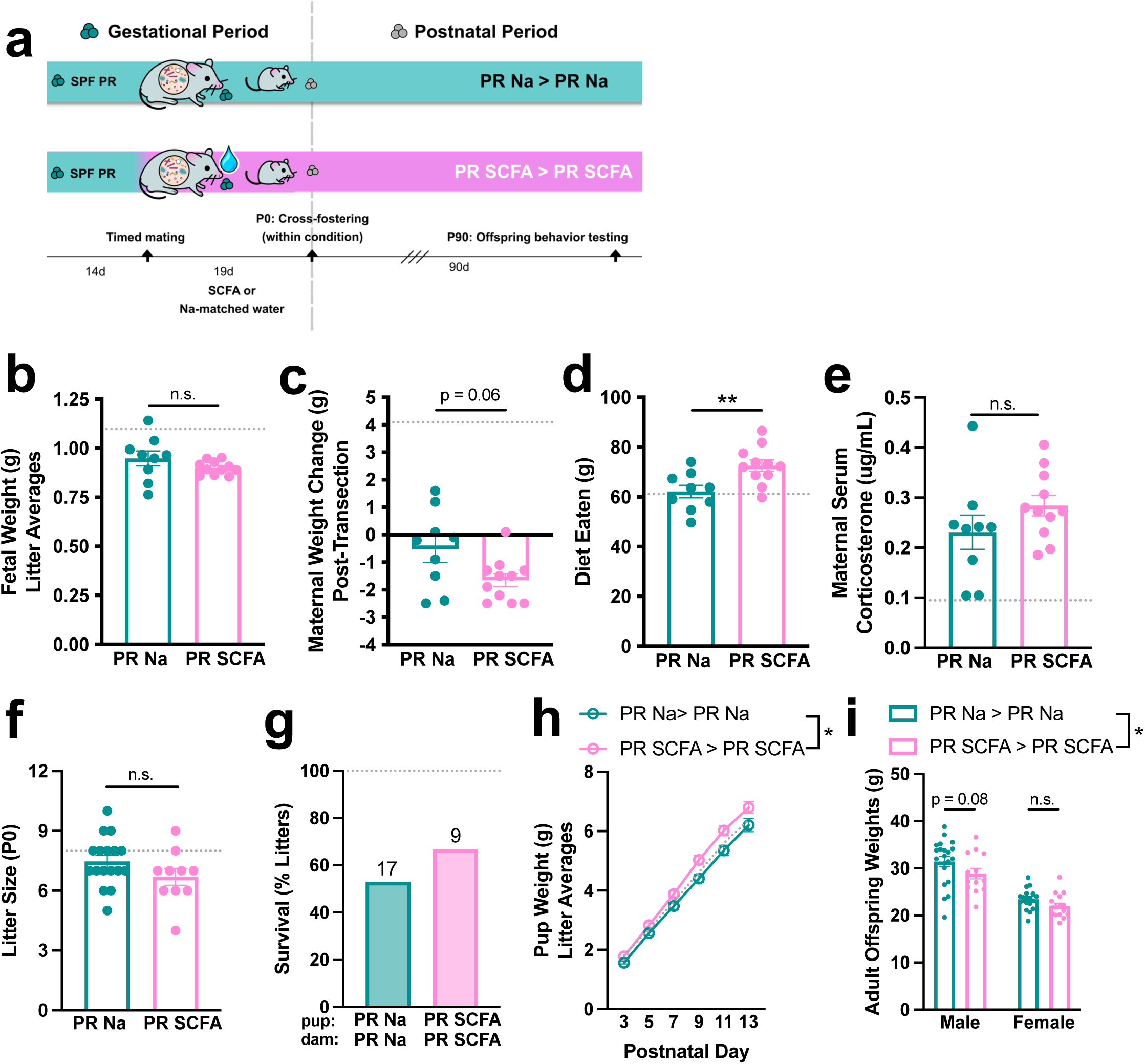
Maternal microbial SCFA supplementation has limited impact on gross measures of maternal and offspring health. **a,** Graphic of gestational supplementation and cross-fostering paradigm for SCFA experiments. **b,** Fetal weight at E18.5 from SPF PR + Na and SPF PR + SCFA dams, all fetuses averaged within each litter (unpaired Welch’s t-test, n = 9, 11, from left to right). Dotted line indicates average value for SPF CD fetuses. **c,** Maternal weight change, from E0.5 to E18.5 post-transection, in SPF PR + Na and SPF PR + SCFA dams (unpaired Welch’s t-test, n = 9, 11, from left to right). Dotted line indicates average value for SPF CD dams. **d,** Diet eaten, from E0.5 to E18.5 in SPF PR + Na and SPF PR + SCFA dams (unpaired Welch’s t-test, n = 9, 11, from left to right). Dotted line indicates average value for SPF CD dams. **e,** Corticosterone measured in serum in SPF PR + Na and SPF PR + SCFA dams at E18.5 (unpaired Welch’s t-test, n = 9, 11, from left to right). Dotted line indicates average value for SPF CD dams. **f,** Litter size, pups per litters, measured at P0, from SPF PR + Na and SPF PR + SCFA dams (unpaired Welch’s t-test, n = 17, 10, from left to right). Dotted line indicates average value for SPF CD litters. **g,** Litter survival (percentage of total litters), from SPF PR + Na pups > SPF PR + Na dams and SPF PR + SCFA pups > SPF PR + SCFA dams (n = 17, 9, from left to right). Dotted line indicates average value for SPF CD litters. Top row refers to pup condition, bottom row refers to dam condition. **h,** Pup weights, all offspring averaged within each litter (two-way repeated measures ANOVA with Sidak, n = 9, 6 litters per group, from top to bottom). Dotted line indicates average value for SPF CD litters. **i,** Adult weights, male and female offspring (two-way ANOVA with Sidak, n = 21, 14, 18, 16, from left to right). Mean +/- SEM, *p < 0.05, **p < 0.01, ***p < 0.001, ****p < 0.0001, n.s. not significant.

**Extended Data Figure 8:**
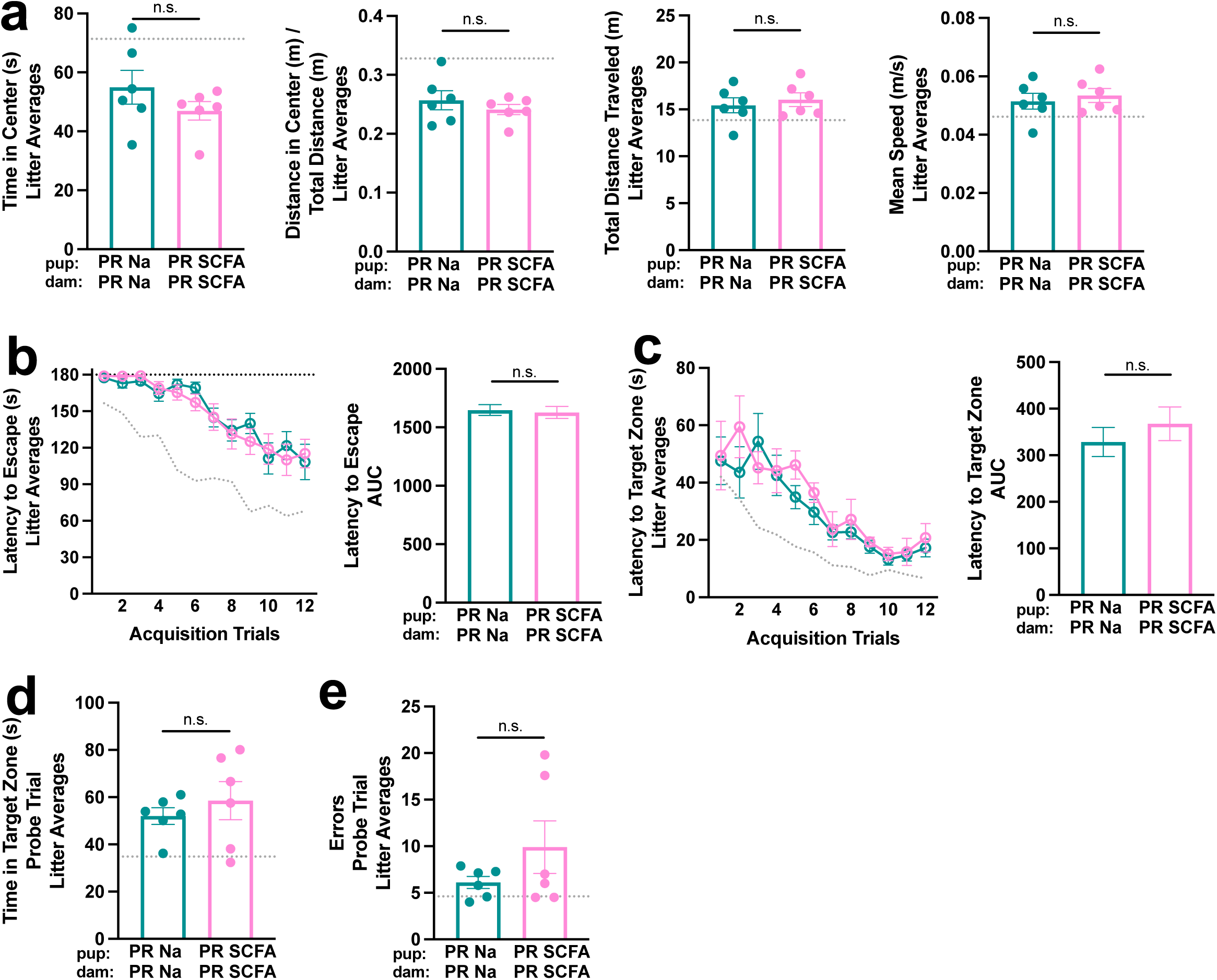
Maternal microbial SCFA supplementation does not influence anxiety-like, locomotor, or cognitive behavioral measures in adult offspring exposed to gestational protein restriction. **a,** Left: Time in center in open field test, all offspring averaged within each litter (Mann Whitney test, n = 6 litters per group). Top row refers to pup condition, bottom row refers to dam condition. Dotted line indicates average value for SPF CD litters. Left-middle: Distance in center in open field test, controlled by total distance traveled, all offspring averaged within each litter (unpaired Welch’s t-test, n = 6 litters per group). Top row refers to pup condition, bottom row refers to dam condition. Dotted line indicates average value for SPF CD litters. Right-middle: Total distance traveled in open field test, all offspring averaged within each litter (unpaired Welch’s t-test, n = 6 litters per group). Top row refers to pup condition, bottom row refers to dam condition. Dotted line indicates average value for SPF CD litters. Right: Mean speed in open field test, all offspring averaged within each litter (unpaired Welch’s t-test, n = 6 litters per group). Top row refers to pup condition, bottom row refers to dam condition. Dotted line indicates average value for SPF CD litters. **b,** Left: Latency to escape in Barnes maze, all offspring averaged within each litter (n = 6 litters per group). Dotted line indicates average value for SPF CD litters. Right: AUC of latency to escape (unpaired Welch’s t-test). Top row refers to pup condition, bottom row refers to dam condition. **c,** Left: Latency to target zone in Barnes maze, all offspring averaged within each litter (n = 6 litters per group). Dotted line indicates average value for SPF CD litters. Right: AUC of latency to target zone (unpaired Welch’s t-test). Top row refers to pup condition, bottom row refers to dam condition. **d,** Time in target zone in Barnes maze probe trial, all offspring averaged within each litter (unpaired Welch’s t-test, n = 6 litters per group). Dotted line indicates average values for SPF CD litters. **e,** Errors made in Barnes maze probe trial, all offspring averaged within each litter (unpaired Welch’s t-test, n = 6 litters per group). Dotted line indicates average values for SPF CD litters. Mean +/- SEM, *p < 0.05, **p < 0.01, ***p < 0.001, ****p < 0.0001, n.s. not significant.

**Extended Data Figure 9:**
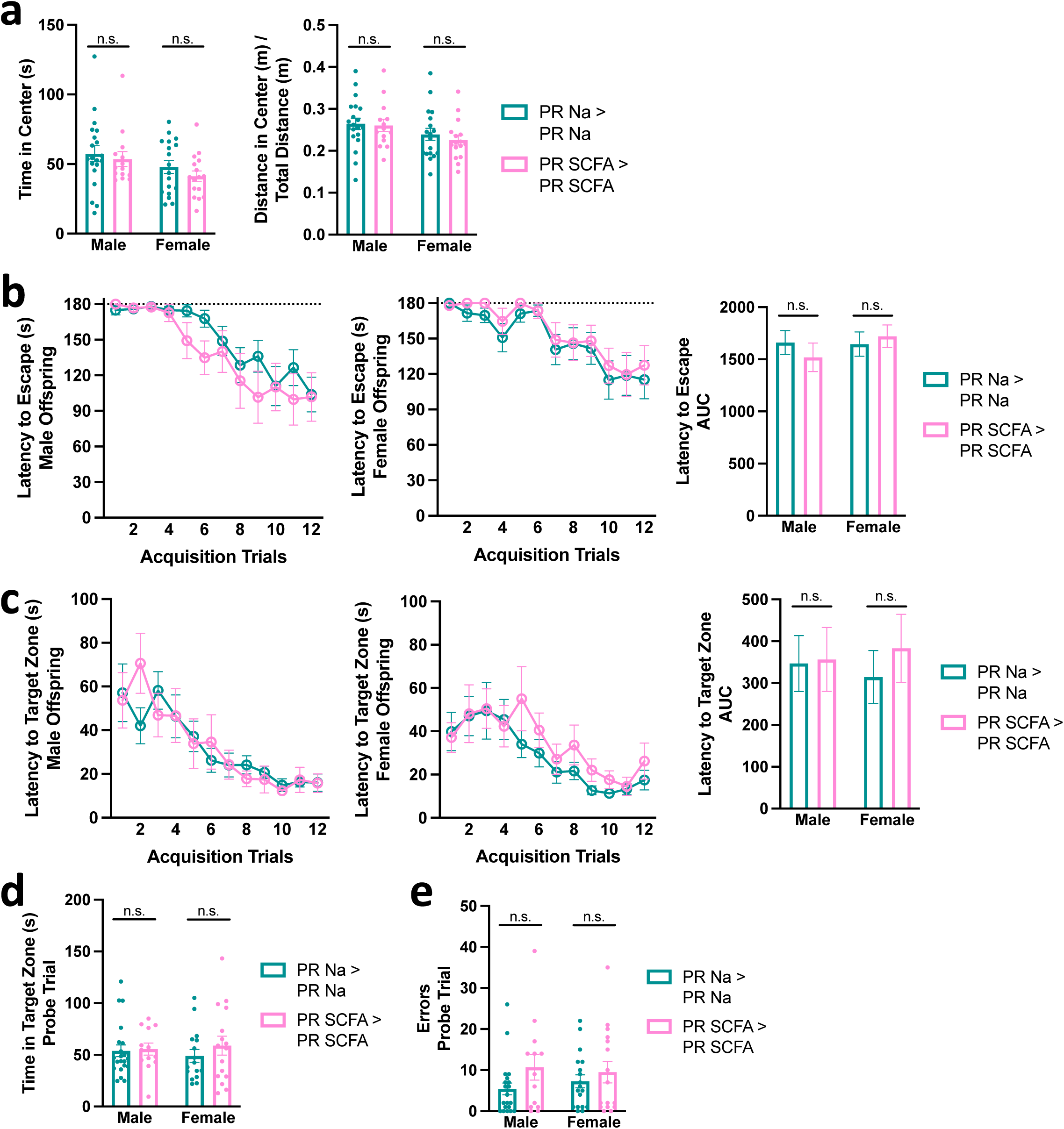
Adult offspring do not exhibit sexually dimorphic behavioral responses to gestational protein restriction and microbial SCFA supplementation. **a,** Left: Time in center in open field test, male and female offspring (Mann Whitney test for each sex, n = 20, 14, 18, 16, from left to right). Right: Distance in center in open field test, controlled by total distance traveled, male and female offspring (unpaired Welch’s t-test for each sex, n = 20, 14, 18, 16, from left to right). **b,** Left: Latency to escape in Barnes maze acquisition phase, male offspring (n = 21, 13). Middle: Latency to escape in Barnes maze acquisition phase, female offspring (n = 18, 16). Right: AUC of latency to escape (unpaired Welch’s t-test for each sex). **c,** Left: Latency to target zone in Barnes maze acquisition phase, male offspring (n = 21, 13). Middle: Latency to target zone in Barnes maze acquisition phase, female offspring (n = 18, 16). Right: AUC of latency to target zone (unpaired Welch’s t-test for each sex). **d,** Time in target zone in Barnes maze probe trial, male and female offspring (Mann-Whitney test for each sex, n = 21, 13, 18, 16, from left to right). **e,** Errors made in Barnes maze probe trial, male and female offspring (Mann-Whitney test for each sex, n = 21, 13, 18, 16, from left to right). Mean +/- SEM, *p < 0.05, **p < 0.01, ***p < 0.001, ****p < 0.0001, n.s. not significant.

**Extended Data Figure 10:**
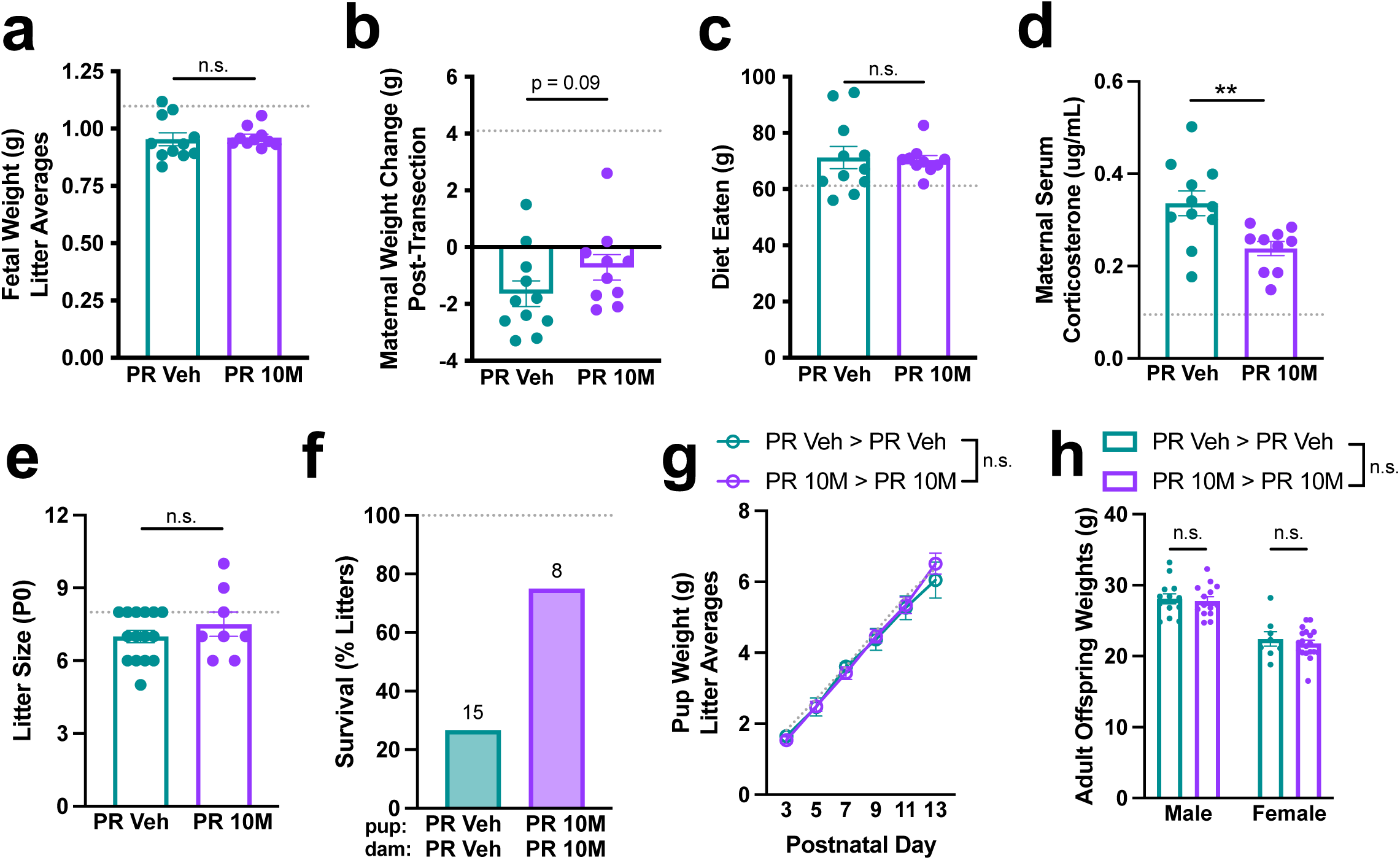
Maternal 10M supplementation has limited impact on gross measures of maternal and offspring health. **a,** Fetal weight at E18.5 from SPF PR + Veh and SPF PR + 10M dams, all fetuses averaged within each litter (unpaired Welch’s t-test, n = 11, 10, from left to right). Dotted line indicates average value for SPF CD fetuses. **b,** Maternal weight change, from E0.5 to E18.5 post-transection, in SPF PR + Veh and SPF PR + 10M dams (Mann-Whitney test, n = 11, 10, from left to right). Dotted line indicates average value for SPF CD dams. **c,** Diet eaten, from E0.5 to E18.5 in SPF PR + Veh and SPF PR + 10M dams (Mann-Whitney test, n = 11, 10, from left to right). Dotted line indicates average value for SPF CD dams. **d,** Corticosterone measured in serum in SPF PR + Veh and SPF PR + 10M dams at E18.5 (unpaired Welch’s t-test, n = 11, 10, from left to right). Dotted line indicates average value for SPF CD dams. **e,** Litter size, pups per litters, measured at P0, from SPF PR + Veh and SPF PR + 10M dams (Mann-Whitney test, n = 15, 8, from left to right). Dotted line indicates average value for SPF CD litters. **f,** Litter survival (percentage of total litters), from SPF PR + Veh pups > SPF PR + Veh dams and SPF PR + 10M pups > SPF PR + 10M dams (n = 15, 8, from left to right). Dotted line indicates average value for SPF CD litters. Top row refers to pup condition, bottom row refers to dam condition. **g,** Pup weights, all offspring averaged within each litter (two-way repeated measures mixed effects analysis with Sidak, n = 4, 6 litters per group, from top to bottom). Dotted line indicates average value for SPF CD litters. **h, h,** Adult weights, male and female offspring (two-way ANOVA with Sidak, n = 13, 14, 8, 19, from left to right). Mean +/- SEM, *p < 0.05, **p < 0.01, ***p < 0.001, ****p < 0.0001, n.s. not significant.

**Extended Data Figure 11:**
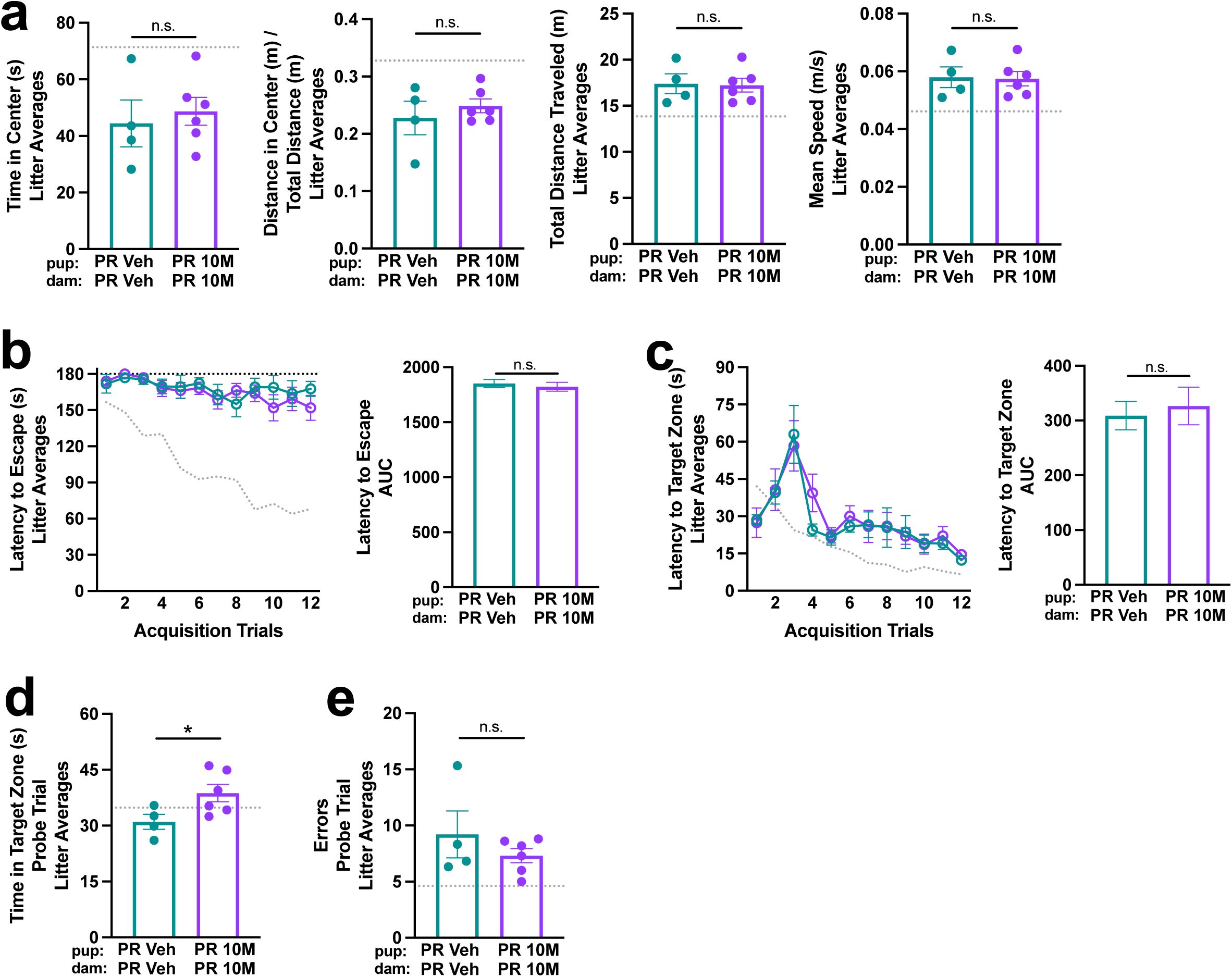
Maternal 10M supplementation does not influence anxiety-like, locomotor, or cognitive behavioral measures in adult offspring exposed to gestational protein restriction. **a,** Left: Time in center in open field test, all offspring averaged within each litter (unpaired Welch’s t-test, n = 4, 6 litters per group, from left to right). Top row refers to pup condition, bottom row refers to dam condition. Dotted line indicates average value for SPF CD litters. Left-middle: Distance in center in open field test, controlled by total distance traveled, all offspring averaged within each litter (unpaired Welch’s t-test, n = 4, 6 litters per group, from left to right). Top row refers to pup condition, bottom row refers to dam condition. Dotted line indicates average value for SPF CD litters. Right-middle: Total distance traveled in open field test, all offspring averaged within each litter (unpaired Welch’s t-test, n = 4, 6 litters per group, from left to right). Top row refers to pup condition, bottom row refers to dam condition. Dotted line indicates average value for SPF CD litters. Right: Mean speed in open field test, all offspring averaged within each litter (unpaired Welch’s t-test, n = 4, 6 litters per group, from left to right). Top row refers to pup condition, bottom row refers to dam condition. Dotted line indicates average value for SPF CD litters. **b,** Left: Latency to escape in Barnes maze, all offspring averaged within each litter (n = 4, 6 litters per group, from top to bottom). Dotted line indicates average value for SPF CD litters. Right: AUC of latency to escape (unpaired Welch’s t-test). Top row refers to pup condition, bottom row refers to dam condition. **c,** Left: Latency to target zone in Barnes maze, all offspring averaged within each litter (n = 4, 6 litters per group, from top to bottom). Dotted line indicates average value for SPF CD litters. Right: AUC of latency to target zone (unpaired Welch’s t-test). Top row refers to pup condition, bottom row refers to dam condition. **d,** Time in target zone in Barnes maze probe trial, all offspring averaged within each litter (unpaired Welch’s t-test, n = 4, 6 litters per group, from left to right). Dotted line indicates average values for SPF CD litters. **e,** Errors made in Barnes maze probe trial, all offspring averaged within each litter (unpaired Welch’s t-test, n = 4, 6 litters per group, from left to right). Dotted line indicates average values for SPF CD litters. Mean +/- SEM, *p < 0.05, **p < 0.01, ***p < 0.001, ****p < 0.0001, n.s. not significant.

**Extended Data Figure 12:**
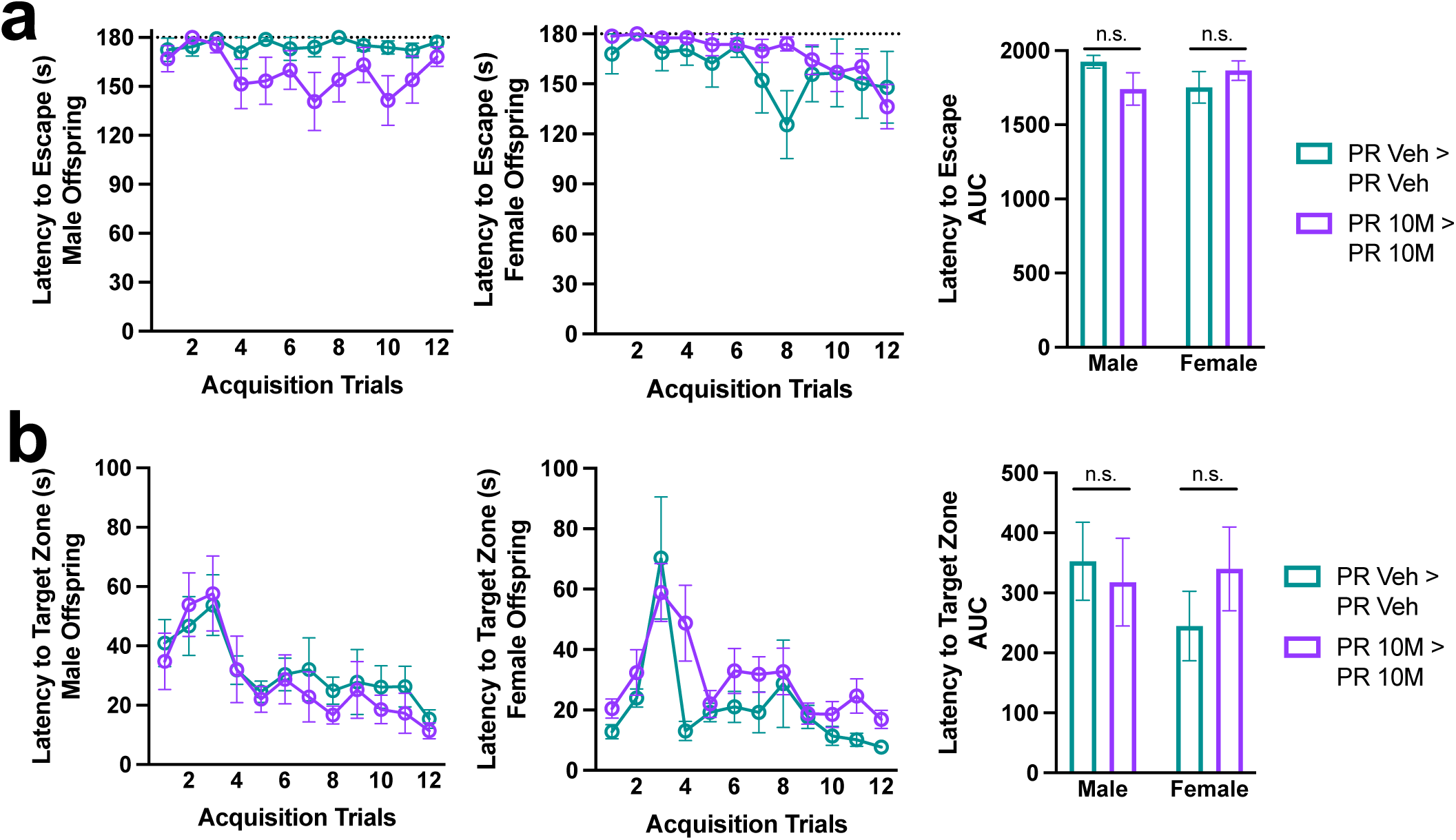
Adult offspring do not exhibit sexually dimorphic behavioral responses to gestational protein restriction and microbial 10M supplementation. **a,** Left: Latency to escape in Barnes maze acquisition phase, male offspring (n = 13, 14). Middle: Latency to escape in Barnes maze acquisition phase, female offspring (n = 9, 18). Right: AUC of latency to escape (unpaired Welch’s t-test for each sex). **b,** Left: Latency to target zone in Barnes maze acquisition phase, male offspring (n = 13, 14). Middle: Latency to target zone in Barnes maze acquisition phase, female offspring (n = 9, 18). Right: AUC of latency to target zone (unpaired Welch’s t-test for each sex). Mean +/- SEM, *p < 0.05, **p < 0.01, ***p < 0.001, ****p < 0.0001, n.s. not significant.

